# AlphaFold3 for Structure-guided Ligand Discovery

**DOI:** 10.64898/2025.12.04.692352

**Authors:** Kartikeya M. Menon, Aakash Davasam, Guo Chen, Claire Bryant, Zifang Deng, Brandon Lam, Chao Yang, Dylan Barcelos, Fangyu Liu, Assaf Alon, Jiankun Lyu

**Author notes:** These authors contributed equally (Kartikeya M. Menon, Aakash Davasam, Guo Chen, Claire Bryant, Zifang Deng).

## Abstract

Deep-learning methods for protein structure prediction, such as AlphaFold2 (AF2) and RosettaFold (RF), have transformed structural biology and accelerated downstream biological discovery. More recent models, including AlphaFold3 (AF3) and RosettaFold All-Atom (RFAA), extend this capability to protein-ligand co-folding, enabling direct prediction of bound complexes from sequence and ligand inputs. This advance has raised the possibility that such models could function not only as structure predictors but also as AI-based molecular docking engines for virtual screening. Yet their true impact on ligand discovery remains unclear and, in many cases, controversial. Here, we systematically assess AF3 co-folding across tasks central to small-molecule discovery compared to conventional molecular docking. First, retrospective enrichment of actives over decoys showed that AF3 outperforms the physics-based docking program DOCK3 across 43 drug targets in DUDE-Z; however, this advantage is largely driven by hidden ligand-only biases inherent to computational decoy sets. In contrast, in three large experimental datasets (the sigma-2 receptor (σ_2_), the D_4_ dopamine receptor (D_4_), AmpC β-lactamase (AmpC)) with over 2,500 tested molecules that lack such biases, DOCK3 achieved stronger overall enrichment, while AF3 contributed mainly to early enrichment. Second, out-of-sample pose reproduction on >8,000 protein-ligand complexes deposited after the AF3 training and validation cutoff showed that AF3 accuracy is strongly dependent on training-set similarity, indicating that the model memorizes atomic positions more than learning general principles of molecular recognition. Finally, in the first prospective head-to-head screen against the σ_2_ receptor, novel to AF3’s training set, AF3 achieved a 13% hit rate and identified a 13 nM binder directly from the screen. However, compared with the parallel DOCK3 campaign, AF3 delivered a two-fold lower hit rate, despite yielding a similar affinity distribution among the top five hits. A crystal structure of the σ_2_ receptor bound to the most potent AF3-derived hit confirmed that AF3 produced a near-native ligand pose. AF3 therefore represents the beginning of deep-learning-based structure-guided ligand discovery: a complementary tool rather than a replacement for conventional docking, with practical applications both as a screening engine and as a post-docking filter that improves hit rates. More broadly, this work establishes a framework for evaluating next-generation deep-learning co-folding models and quantifying their impact on small-molecule discovery.

## Introduction

Deep-learning-based protein structure prediction has revolutionized structural biology and protein design, most notably through AlphaFold2 (AF2)^1^ and RosettaFold (RF)^2^, which deliver near-experimental accuracy for thousands of proteins. This transformative contribution was recognized with the 2024 Nobel Prize in Chemistry. Beyond individual cases, the impact of AF2 has been felt at scale: the AlphaFold database now provides structural models for the entire human proteome and for 47 additional organisms, encompassing more than 200 million proteins and nearly all therapeutically relevant targets^3–5^. These predicted structures have already proven invaluable across a wide range of applications, including protein-protein interaction mapping^6–8^, target and function prediction^9,10^, and elucidating biological mechanisms of action^11–13^.

A natural next question is whether such predicted structures can also accelerate ligand discovery. Retrospective docking studies suggest that AF2 models struggle to recapitulate ligand binding modes and to distinguish actives from decoys in ligand discovery benchmarks when compared with experimental structures^14–24^. However, retrospective analyses do not address whether AF2-predicted structures prospectively guide ligand discovery. Prospective testing has provided more encouraging results^25,26^. Docking a 16-million compound library to AF2-predicted models of the trace amine-associated receptor 1 (TAAR1) outperformed homology models, producing a nearly three-fold higher hit rate (60% vs. 22%) and yielding the most potent agonists, including compounds with in vivo efficacy in antipsychotic-like behavioral assays. These results demonstrated that AF2-predicted structures serve as effective templates for virtual screening, directly enabling therapeutic discovery. Complementary results emerged from head-to-head comparisons between experimental and AF2-predicted structures for the σ_2_ and 5-HT2A receptors, using ultra-large make-on-demand libraries (490 million for σ_2_ and 1.6 billion for 5-HT2A). Strikingly, docking to AF2-predicted models yielded results comparable to those obtained with experimental structures. Docking to the σ_2_ AF2 model achieved a 54% hit rate versus 51% for the crystal structure, with best binders of 1.6 nM and 1.8 nM, respectively. Similarly, AF2 models of 5-HT2A delivered a 26% hit rate, comparable to the 23% achieved with the cryo-EM structure, and identified both agonists and antagonists with nanomolar to micromolar potencies. These prospective examples establish AF2 models as practical substitutes for experimental structures in ligand discovery campaigns, potentially accelerating projects by years when no experimental structure is available.

Building on AF2, recent diffusion-based approaches^27–31^ extend prediction capabilities to nucleic acids, ions, and small molecules, resulting in co-folding models, such as AlphaFold3 (AF3)^32^ and others^33–39^. Importantly, these methods enable direct protein–ligand co-folding, generating complex structures without the need for subsequent docking. With AF3, this not only expands the spectrum of biological structure predictions^40^ but also raises the possibility of using AF3 itself as a docking engine for screening campaigns. However, their true impact on ligand discovery remains unclear. Before co-folding methods emerged, several retrospective studies reported that deep-learning (DL)-based docking methods failed to consistently outperform molecular docking when evaluated under both physical plausibility and pose reproduction metrics outside their training distributions.^41,42^ More recent studies similarly suggest that co-folding approaches, including AF3, rely heavily on memorization of training complexes rather than truly learning principles of molecular interactions^43–51^, limiting their generalizability for novel ligand discovery. Even under extreme binding site abuse (removal, inversion, and packing with phenylalanine residues), AF3 still predicts ligand poses relatively unchanged versus the wildtype binding site..^48^ Within a recent covalent ligand discovery domain, AF3 achieved near-perfect enrichment of covalent ligands over property-matched decoys across nine kinase targets. In a prospective covalent screen against Bruton’s tyrosine kinase (BTK), AF3 also identified a potent and selective inhibitor, whose binding mode was confirmed by co-crystallography with sub-angstrom accuracy.^52^ However, prospective testing of AF3 for non-covalent ligand discovery against a novel target outside its training set has not yet been reported, which is the more common scenario for drug discovery campaigns.

Here we aim to evaluate AF3 co-folding as a practical docking engine for both retrospective and prospective ligand discovery. Molecular docking is generally assessed by two key criteria: enrichment, the ability to prioritize experimentally confirmed actives over inactives, and pose reproduction, the ability to recover experimentally observed ligand–protein binding modes as a measure of structural fidelity. To address these goals, we benchmarked AF3 across four tasks: (1) In-sample pose reproduction: Evaluated on the widely used docking benchmark set DUDE-Z (Directory of Useful Decoys, Enhanced and optimiZed version)^53^, using experimental complexes that are deposited before AF3’s training cutoff to determine its accuracy in recovering bound ligand poses. (2) Retrospective enrichment: tested on DUDE-Z and extended to large experimental compound sets for σ_2_, D_4_, and AmpC from previous ultra-large-scale docking campaigns. (3) Out-of-sample pose reproduction: assessed on a newly curated set of around 10,000 ligand–protein complexes, focusing on out-of-distribution subsets defined by chemical novelty to evaluate generalization. (4) Prospective enrichment: applied in a structure-based ligand discovery campaign against the σ_2_ receptor to assess their ability to drive prospective ligand discovery. Approximately 700,000 monocations from the Enamine in-stock library were screened using both AF3 and the physics-based molecular docking program DOCK3. Finally, we discuss the strengths and limitations of AF3 co-folding in light of these analyses.

## Results

### Retrospective Benchmarking with the DUDE-Z Set

We began our evaluation of AF3 on the DUDE-Z benchmark, a widely used dataset for assessing physics-based molecular docking performance. For each of its 43 protein targets, DUDE-Z provides a ligand-bound experimental structure to template rigid-receptor docking. In addition, each target includes a set of experimental actives (true positives) together with a larger set of property- and charge-matched decoys (false positives), curated at a ratio of roughly 1:50. These decoys closely resemble the actives in physicochemical properties but differ topologically.

Although all DUDE-Z targets appear in the AF3 training set, this benchmark still provides a useful sanity check for AF3’s ability to reproduce ligand-bound experimental poses. As a first step, we evaluated in-sample pose reproduction—testing whether AF3 can recover the crystallographic ligand poses from their corresponding templates. Establishing this baseline is essential, because without reliable pose recovery, any apparent enrichment of actives over decoys would be difficult to interpret. To bypass multiple sequence alignment, we provided AF3 with the experimental protein-ligand structure, thus accelerating AF3 predictions by an order of magnitude (from 20 minutes to 2 minutes) and making it computationally practical to predict thousands of compounds. As expected, all targets using template-based AF3 co-folding achieved pocket RMSDs < 2 Å. Pocket RMSD was calculated using the APoc package^54^, defined by the positions of C-alpha and C-beta atoms of residues 5Å around the ligand. This initial benchmark confirmed that AF3 accurately predicts the protein binding pocket. The more stringent test lies in ligand pose reproduction (blind docking). In this setting, AF3 successfully reproduced experimental poses in 86% (37/43) of DUDE-Z targets, where success is defined by ligand RMSDs below 2 Å (**Fig. 1a**). Because all DUDE-Z targets are included in AF3’s training set, this analysis represents an in-sample upper bound of AF3’s pose reproduction ability. These results demonstrate that template-based AF3 co-folding can reliably recover binding geometries under in-sample conditions, providing a strong structural foundation for subsequent enrichment analyses on DUDE-Z.

**Figure 1.**
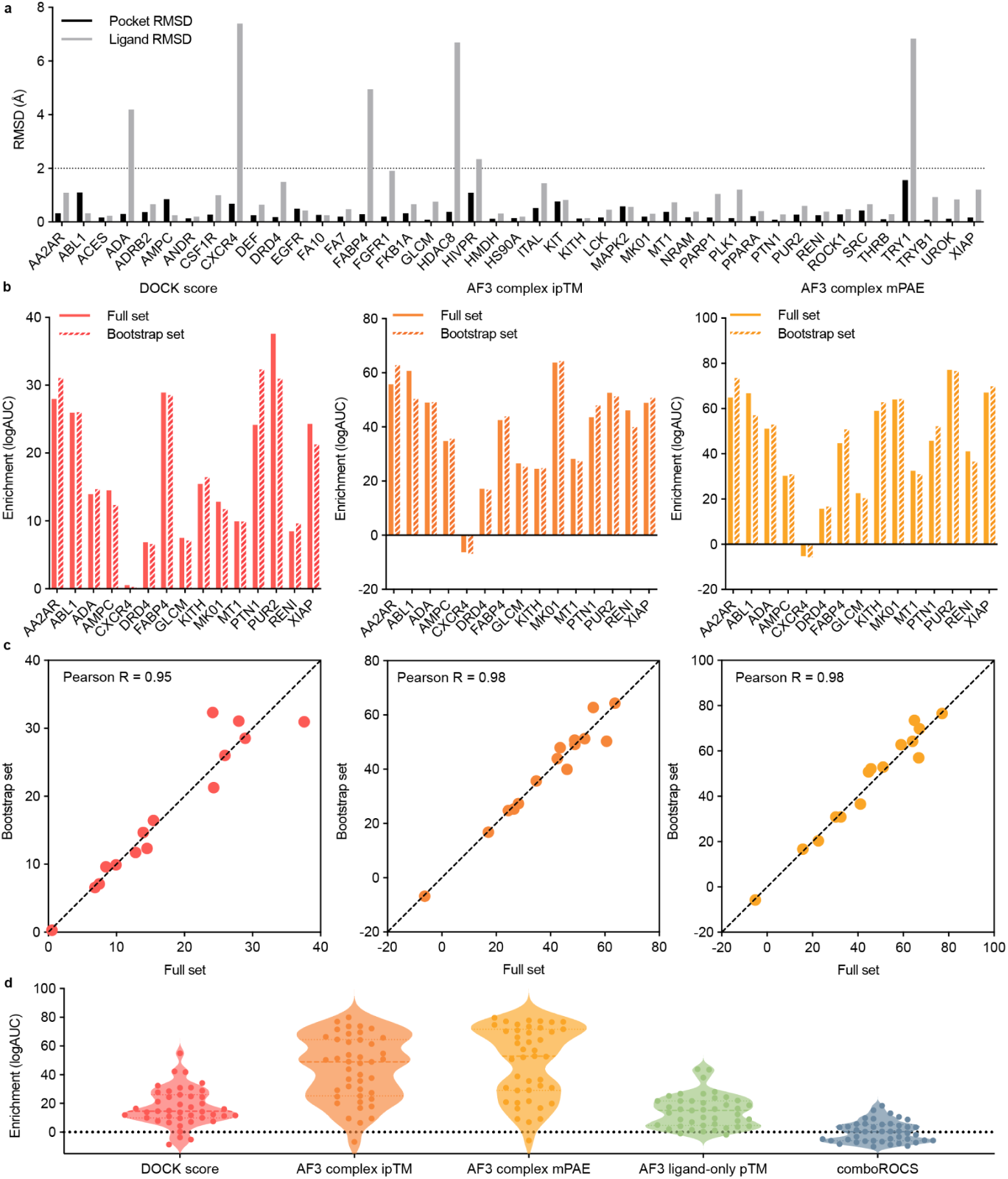
Retrospective benchmarking of template-based AF3 co-folding versus physics-based docking on the DUDE-Z set. **a**. Pose-reproduction accuracy of AF3 across 43 DUDE-Z protein-ligand complexes. Bars show root-mean-square deviations (RMSDs) for binding-pocket residues (black) and ligand poses (gray) relative to experimental structures; the dashed line marks the 2 Å success threshold. **b**. Comparison of enrichment (logAUC) between DOCK3 docking scores and AF3 confidence metrics (ipTM and mPAE) using the complete DUDE-Z ligand/decoy sets (solid bars) and the down-sampled bootstrap subset (hatched bars; 50 actives + 500 decoys per target). **c**. Correlation of enrichment between the full and bootstrap sets for DOCK3 (left), AF3 ipTM (center), and AF3 mPAE (right). **d**. Distribution of target-wise enrichment (logAUC) across five scoring schemes: DOCK3 docking scores, AF3 complex ipTM, AF3 complex mPAE, AF3 ligand-only pTM, and ComboROCS 3D ligand-shape similarity to AF3’s training set. Each point represents one DUDE-Z target; violin plots show the overall distribution across targets. The dashed horizontal line denotes the expected logAUC for a random ranking.

Having established that AF3 can reliably reproduce experimental poses under in-sample conditions, we next asked whether this structural fidelity also improves enrichment in the DUDE-Z benchmark. For each target, actives and decoys were ranked independently by three scoring functions: AF3 ipTM (interface predicted Template Modeling score: higher values indicate greater confidence in the protein-ligand interface), AF3 mPAE (mean Predicted Aligned Error: lower values reflect lower model uncertainty), and the DOCK3 score (physics-based energy score: kcal/mol). Enrichment was quantified using both logAUC (area under the semi-log adjusted ROC curve, emphasizing early enrichment) and AUC (standard ROC area). Due to the high computing cost of running AF3 predictions, we ran 15 DUDE-Z systems with a full set of actives and decoys, totaling 77,396 molecules (1,482 actives), representing 18.0% of the total DUDE-Z library (n=430,141). For these 15 targets, the logAUC and AUC between the full and bootstrapped sets correlates well with a Pearson correlation coefficient (*r*) of 0.95-0.98 and of 0.98-0.99 for among three scoring metrics, respectively (**Fig. 1b** and **1c**; **Extended Data Fig. 1a** and **1b**), justifying the usage of the bootstrapped set to evaluate the performance difference between these two docking methods. On the bootstrapped data set (50 ligands and 500 charge- and property-matched decoys), AF3 metrics achieve improved enrichment over DOCK3 score in 93% (40/43) of all DUDE-Z targets with an average logAUC improvement of 27 and 31, respectively (**Fig. 1d**; **Extended Data Fig. 2a, 2b** and **2c**). We compared enrichment with ipTM and mPAE and found comparable results. Among the four targets where AF3 yielded worse enrichment, two (CXCR4 and TRY1) also demonstrated poor ligand RMSD performance (**Fig. 1a**), suggesting that inaccurate ligand placement may underlie some of the observed weaknesses in enrichment. To evaluate whether AF3 enrichment performance depends on the physical plausibility of its predicted ligand complexes, we applied PoseBusters (PB), a recently developed validation framework that filters out ligand–protein complexes with chemically or structurally implausible geometries (e.g., strained conformers, violated stereochemistry, protein-ligand clashes).^41^ PoseBusters filtering provides an additional quality control step to ensure that enrichment trends are not driven by artifacts in pose generation because AF3 was not explicitly trained to satisfy such physics constraints. After restricting to only PoseBusters-validated molecules for each DUDE-Z target (**Extended Data Fig. 3a** and **3b**), enrichment values remained strongly correlated with those obtained from the full pared-down datasets. The Pearson correlations between full vs. PB-filtered logAUC values ranged from 0.91 to 0.99 across all five scoring metrics (ComboROCS, DOCK score, ipTM, mPAE, and ligand-only pTM; **Extended Data Fig. 3c**). The logAUC correlations were similarly high, ranging from 0.95 to 0.97 (**Extended Data Fig. 3d**), indicating that structural filtering does not largely affect ligand-decoy ranking performance. Consistent with the unfiltered analysis, the enrichment trends driven by AF3 scores (ipTM and mPAE) continued to outperform DOCK3 for the vast majority of targets after PB filtering. These results confirm that the observed enrichment advantage of AF3 is retained when considering only physically plausible binding poses. Although AF3 was not originally trained to distinguish actives and inactives through structural inference, the performance across 43 DUDE-Z systems suggest that AF3 metric scores are capable of this classification.

**Figure 2.**
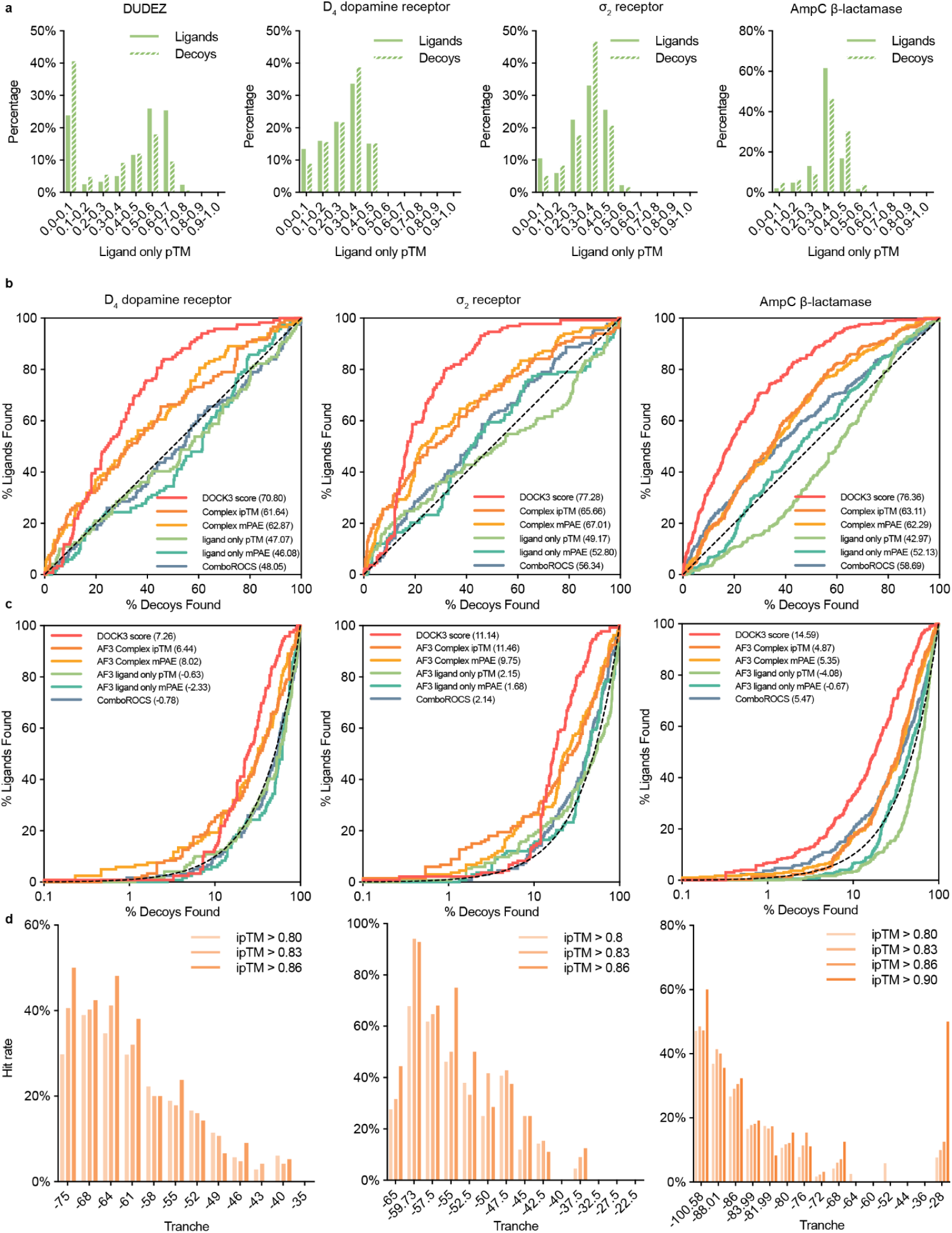
Benchmarking AF3 and DOCK3 enrichment across DUDE-Z and large experimental test sets. **a.** Distribution of AF3 ligand-only pTM scores for actives (solid) and decoys (hatched) in the DUDE-Z benchmark and three experimental datasets: D_4_ dopamine receptor, σ_2_ receptor, and AmpC β-lactamase. **b.** ROC curves comparing enrichment of actives over decoys for D_4_ (left), σ_2_ (center), and AmpC (right) datasets. **c.** Log adjusted ROC plots for the same datasets in **b**. **d**. Experimental hit-rate analysis for D_4_, σ_2_, and AmpC docking campaigns. Hit rate is calculated as the number of hits divided by the number of tested per docking-score tranche.

**Figure 3.**
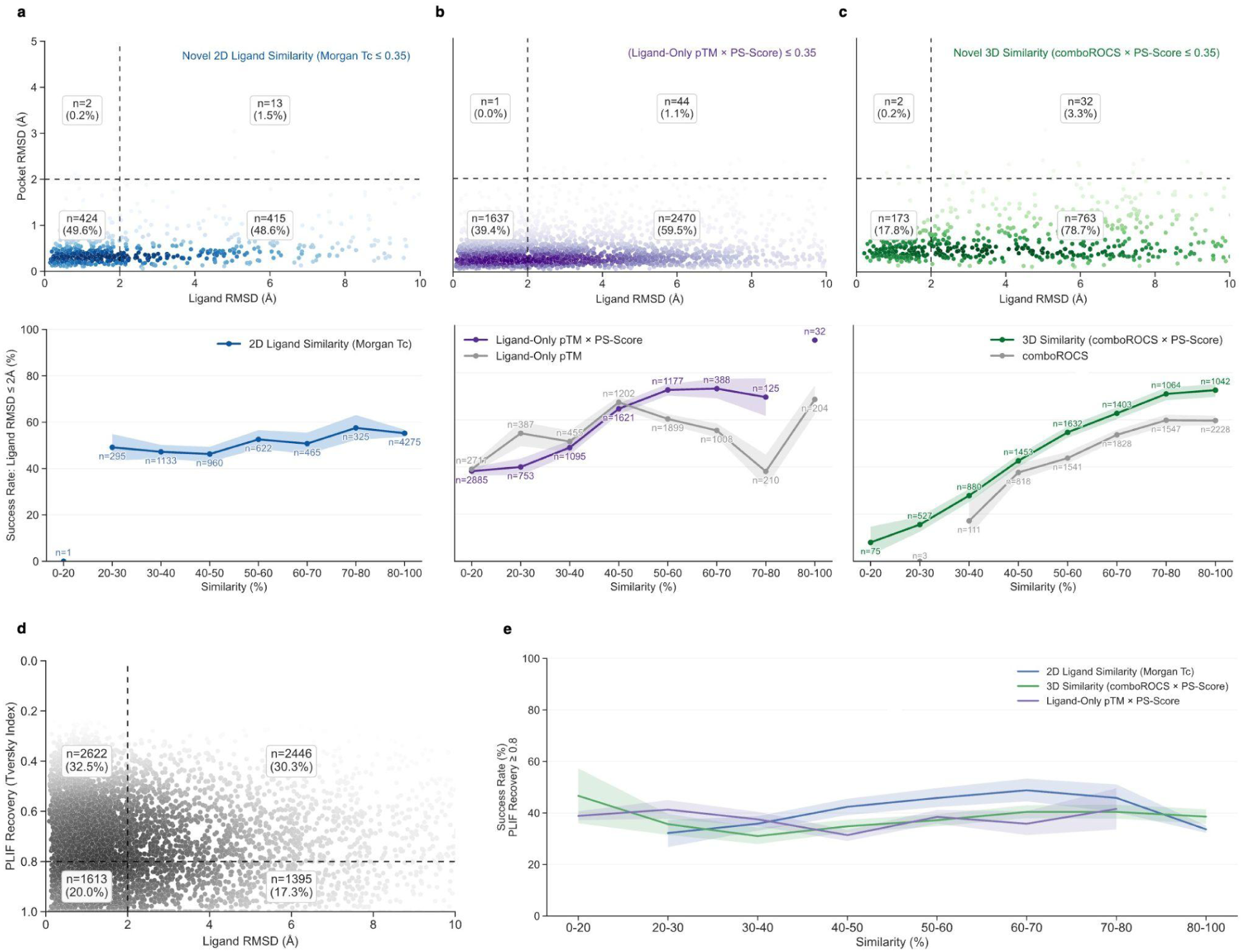
Out-of-sample generalization and memorization analysis of AF3 co-folding across novel protein–ligand complexes. **a–c (top panels).** Ligand RMSD versus pocket RMSD for AF3-predicted complexes under three definitions of novelty relative to the AF3 training set: **a.** 2D ligand chemical novelty (Morgan Tc ≤ 0.35); **b.** novelty based on ligand-only pTM × pocket similarity (PS-Score ≤ 0.35); **c.** novelty based on 3D ligand–pocket similarity (comboROCS × PS-Score ≤ 0.35). Dashed lines indicate the 2 Å success threshold for ligand and pocket RMSD, and annotated regions show the fraction of complexes falling into each quadrant. Quadrant interpretation: upper-right = incorrect ligand & incorrect pocket; upper-left = correct pocket but incorrect ligand; lower-right = correct ligand but incorrect pocket; lower-left = correct ligand & correct pocket (full success). **a–c (bottom panels).** Success rate of accurate ligand pose reproduction (ligand RMSD ≤ 2 Å) across similarity bins for the same three novelty metrics: **a.** 2D Morgan Tc; **b.** ligand-only pTM (gray) and ligand-only pTM × PS-Score (purple); **c.** comboROCS × PS-Score (green) compared with comboROCS alone (gray). Shaded areas correspond to the 95% confidence interval for each bin, calculated from 1,000 bootstrap samples. **d.** Protein–ligand interaction fingerprint (PLIF) recovery (inverted y-axis) versus ligand RMSD for all complexes, showing that low ligand RMSD does not guarantee recovery of native interaction patterns. **e.** Success rate of PLIF recovery (Tc ≥ 0.5) across similarity bins for the three metrics in **a–c**, demonstrating that higher similarity to the AF3 training set improves positional accuracy more than interaction recovery. Bins with fewer than 50 observations (n < 50) are displayed as single points without error estimates and line trends, as the sample size is insufficient to reliably calculate uncertainty.

Given AF3’s strong enrichment performance on DUDE-Z, we next asked whether this signal truly arose from protein–ligand structural prediction or instead explained by hidden biases of the dataset. Previous work has shown that property-matched decoy-based benchmarks such as DUDE and its DUDE-Z contain latent ligand-level artifacts that can mislead deep learning models into classifying actives and decoys without relying on protein context.^55,56^ To probe this possibility, we tested a ligand-only pTM score, in which AF3 was run using only the ligand SMILES string as input, without providing either the protein sequence or an experimental structure template. This setting removes protein information entirely, so any enrichment observed must originate from biases in ligand chemistry rather than recognition of protein–ligand interactions. As shown in **Fig. 1d**, AF3 ligand-only pTM still produced non-random enrichment in 84% (36/43) of DUDE-Z targets, with logAUC and AUC values well above random expectation. In several cases, ligand-only enrichment approached the performance of DOCK3 or full AF3 scoring, suggesting that DUDE-Z’s actives and decoys can often be separated based on ligand features alone. To further quantify the extent of ligand-only bias in DUDE-Z, we directly compared enrichment using AF3 ligand-only pTM versus DOCK3 docking scores. As shown in **Extended Data Fig. 2d**, 47% (20/43) of targets achieved comparable or better enrichment with ligand-only pTM relative to DOCK3, with ΔlogAUC values at or above zero. These findings underscore that while AF3 can achieve superior enrichment on DUDE-Z, a substantial component of the signal may reflect ligand-only bias. This result reinforces the importance of moving beyond decoy-based benchmarks and toward prospective or out-of-distribution tests to more accurately measure AF3’s capability for structure-based virtual screening.

To further test whether AF3’s superior enrichment on DUDE-Z is driven by memorization of its training data, we compared its performance against ComboROCS from OpenEye’s Rapid Overlay of Chemical Structures (ROCS). ComboROCS quantifies 3D Tanimoto similarity by measuring the overlap of molecular volume (“shape”) and chemical features (“color”) between two ligand poses^57^. The AF3 training set was curated from protein-ligand complexes deposited in the RCSB database prior to September 2021, the final dataset contained 30,528 unique ligand pose entries (See **Methods** for more details). For each DUDE-Z molecule, we calculated its ComboROCS similarity to every ligand pose in the AF3 training set and retained the highest score as its maximum training-set similarity. Enrichments (AUC or logAUC) were then evaluated based on these ligand-only pTM and ComboROCS similarity scores. As shown in **Extended Data Fig. 2e**, 88% (38/43) of DUDE-Z targets showed comparable or better enrichment using AF3 ligand-only pTM versus ComboROCS similarity. In many cases, ligand-only AF3 outperformed ComboROCS, suggesting that 3D shape and pharmacophore similarity to the AF3 training set is not sufficient to discriminate between actives and decoys. Together, these results suggest that the superior enrichment of AF3 ipTM or mPAE scoring is not due to memorization of specific ligand poses. Instead, as shown in **Extended Data Fig. 4**, AF3’s ligand-only single and pairwise embeddings exhibit broad pTM gradients, with actives more frequently occupying higher-pTM regions. This indicates that the bias arises from nuanced AF3 ligand representations rather than rigid 3D shape or pharmacophore similarity to its training set.

To summarize this section, we demonstrate that while AF3 consistently achieves superior enrichment over physics-based docking on the DUDE-Z benchmark, much of this advantage can be attributed to hidden biases inherent in property- and charge-matched decoy datasets. Ligand-only pTM scoring metrics reproduce a large fraction of the enrichment signal, indicating that performance on DUDE-Z does not necessarily reflect true protein-ligand recognition. Thus, DUDE-Z should be regarded as an early validation for deep learning–based approaches, confirming their ability to recover poses and distinguish actives from decoys under favorable conditions. However, DUDE-Z should not be taken as a realistic benchmark for retrospective virtual screening. The fact that AF3 and related co-folding methods always outperform physics-based docking on DUDE-Z highlights the limitations of such computationally generated decoy datasets. This, in turn, motivates the question of what constitutes an appropriate benchmark for testing AF3 and other structure-based deep learning approaches in real-world discovery settings.

### Benchmark on large experimental testing sets from ultra-large library docking campaigns against AMPC, D_4_, and σ_2_

Given that AF3’s superior performance on DUDE-Z largely reflects hidden biases, the next question is whether such biases persist in more realistic experimental benchmarks. To test this, we examined the distribution of ligand-only pTM scores for actives and inactives across DUDE-Z and three experimental data sets from ultra-large library docking campaigns against D_4_, σ_2_, and AmpC, each containing at least 400 tested compounds with diverse scaffolds. As expected, DUDE-Z actives were skewed toward higher ligand-only pTM bins relative to decoys. In contrast, the experimental test sets displayed nearly overlapping distributions between actives and inactives, indicating that ligand-only pTM scores are not sufficient to discriminate between the two groups (**Fig. 2a**). Consistent with this, t-SNE projections of AF3 ligand-only embeddings across all four datasets reveal that DUDE-Z actives concentrate in islands of high predicted pTM, whereas compounds from the experimental sets (actives and inactives) are intermingled and lack such high-pTM-enriched regions (**Extended Data Fig. 4**). This analysis confirms that ligand-only biases are specific to computationally generated decoy libraries such as DUDE-Z and not within prospective make-on-demand screening sets. Additionally, we previously showed that molecules from ultra-large virtual libraries tend to have low similarity to bio-like molecules, including worldwide drugs, metabolites, and natural products.^58^ Because AF3’s training set is derived from the PDB, where structural biologists have historically focused on complexes with such bio-like compounds due to their biological or therapeutic importance, ultra-large library molecules are less likely to resemble training set ligands and are therefore less prone to memorization bias. For the three experimental testing sets (σ_2_, D_4_, and AmpC)^14,59,60^, ligand-only AF3 metrics (pTM and mPAE) behave randomly. The logAUC values are close to or below zero across all targets (**Fig. 2b** and **2c**; D_4_: -0.63 or -2.33, σ_2_: 2.15 or 1.68, AmpC: -4.08 or -0.67), confirming that these datasets are free of the ligand-only bias in DUDE-Z and thus serve as more realistic benchmarks. Similarly, ComboROCS displayed random performance for D_4_ (logAUC: -0.78, AUC: 48.05) and σ_2_ (logAUC: 2.4, AUC: 56.34). Intriguingly, AMPC showed non-random enrichment (logAUC: 5.47, AUC: 58.69), suggesting that the AmpC test set retains some 3D similarity to AF3’s training set and is more susceptible to memorization (**Fig. 2b** and **2c**). When evaluated by overall enrichment (AUC), physics-based docking DOCK3 consistently outperformed AF3 across all three large experimental datasets (**Fig. 2c**). For example, on σ_2_, DOCK3 reached an AUC of 77.28, compared with 65.66 (ipTM) and 67.01 (mPAE), with similarly large gaps on D_4_ and AmpC. In contrast, AF3 improved early enrichment, which is often the relevant regime for virtual screening. Early enrichment was quantified using logAUC and EF_10_, where EF_10_ (enrichment factor at the top 10%) quantifies actives recovered in the top 10% of a ranked library relative to random selection. For σ_2_ and D_4_, ipTM and mPAE exceeded DOCK3 in both logAUC and EF_10_ (**Fig. 2c** and **Extended Data Fig. 5**). However for AmpC, docking improved all enrichment metrics, including early enrichment. Taken together, these results suggest that AF3 confidence scores have potential as rescoring filters applied after docking.

A longstanding question in docking is how well rank predicts binding likelihood. Traditionally, only the very top-ranked molecules are tested, but with the advent of ultra-large make-on-demand libraries, it has become possible to systematically sample across a wide range of docking scores to experimentally define ‘hit-rate’ curves. Here, we define hit rate as the number of experimental hits divided by the total number of tested compounds within a given score bin. This strategy was first applied in the original D_4_ campaign, where 549 molecules were sampled across 12 docking score bins spanning high-, mid-, and low-scoring ranges, yielding a quantitative response curve that related docking score to experimental activity.^59^ AmpC^60^ and σ_2_^14^ datasets were generated in a similar fashion, using binning schemes that enabled statistically meaningful estimates of hit rate as a function of docking score. For the D_4_ dopamine receptor and σ_2_ receptor, the hit rate vs. docking score relationship was sigmoidal, with an early plateau, then a nearly monotonic decline as scores worsened. By contrast, for AmpC β-lactamase, the curve dropped monotonically without a plateau, suggesting continuous access to new ligands as larger libraries were tested.

Using this same binning approach, we re-analyzed the D_4_, σ_2_, and AmpC datasets to assess whether AF3 complex scoring metric, measured by ipTM, could improve experimental hit rates when applied as a post-docking filter. As shown in **Fig. 2d**, applying progressively stricter ipTM thresholds (e.g., > 0.80, > 0.83, > 0.86, or > 0.90, depending on the receptor) consistently increased the hit rate within each docking-score tranche. This effect is most pronounced in the top-ranked bins—where prospective campaigns typically select compounds for testing, but where scoring artifacts are also most common—because the ipTM filter selectively removes false positives that were highly ranked by docking alone. Across all three datasets, applying an ipTM filter substantially improved hit rates in the highest-scoring tranche: for σ_2_, the hit rate increased from 34 % to 68 % with ipTM > 0.83 (2.0-fold); for D_4_, from 39 % to 66 % with ipTM > 0.83 (1.7-fold); and for AmpC, from 26 % to 47 % with ipTM > 0.80 (1.8-fold). Notably, AF3 confidence metrics on their own didn’t correlate with binding affinity, underscoring that their value lies in improving hit rate, not predicting affinity (**Extended Data Fig. 6**). These gains were concentrated in the top docking bins, indicating that ipTM functions as a false-positive filter, enriching the most promising candidates for testing.

The strong enrichment of experimental actives over decoys is supported by the high success rate of ligand pose reproduction. For two of the three large testing set targets (AmpC and σ_2_), nine crystal structures were deposited in the PDB after AF3’s training cutoff but still within its validation window (May 2022 to Jan 2023). Of these nine complexes, AF3 successfully reproduced the ligand pose (ligand RMSD < 2 Å) in four cases. (7MFI: cholesterol in σ_2_, 7M96: Z4857158944 in σ_2_, 9C84: Z6615020275 in AMPC, 9C6P: Z6615017509 in AMPC) versus 6 successful predictions by DOCK3 (7M93: PB-28 in σ_2_, 7M95: Z1241145220 in σ_2_, 7M96, 9DHL: Z6615017782 in AMPC, 9C84, and 9C6P). In the complex prediction of PB-28 and the σ_2_ receptor (AF3-ligand RMSD: 11.9Å, DOCK3-ligand RMSD: 1.21Å), the AF3 pose is rotated 180 degrees versus the docking and crystal poses, creating a hydrogen bond between the ether group of PB-28 and Q77 as opposed to the salt bridge between the ligand’s protonated amine and D29. In the complex prediction of A1AU1 and AMPC β-lactamase, both AF3 (RMSD: 5.66Å) and DOCK3 (RMSD: 5.61Å) reverse the pose against the AMPC reference crystal structure, forming the same hydrogen bond network with S64, A318, Q120, and N152 (**Extended Data Fig. 7**).

### Out-of-Sample Performance

Accurately predicting ligand binding poses is foundational to structure-based drug design, particularly when the aim is to identify truly novel chemotypes rather than close analogs of known compounds. Even for targets that have been previously characterized, the ability to model complexes with ligands that are structurally dissimilar from those seen during training is essential for practical drug discovery applications—ranging from relatively straightforward patent busting to the more demanding task of identifying novel molecular scaffolds. For these reasons, we first evaluated AF3’s ability to generalize at the ligand level, evaluating whether it can correctly model protein-ligand complexes involving ligands that are chemically distinct from those present in its training data.

To rigorously test AF3’s ability to generalize, we curated a benchmark test set of protein-ligand complexes containing chemically novel ligands. Novelty was defined by enforcing a maximum Tanimoto coefficient (Tc) of 0.35 between each test-set ligand (from PDB entries deposited between Jan 2023 and Dec 2024) and all ligands in AF3’s training data (deposited before Sep 2021). This novelty Tc threshold as a way of classifying 2D fingerprint similarity has been commonly used in previous campaigns, and we adopted the same criterion here.^61^

For each complex, we generated 25 structures (5 samples per 5 seeds) and selected the top-ranked model for comparison against its corresponding experimental structure obtained from the BioLip2 database (https://zhanggroup.org/BioLiP) (**Extended Data Fig. 8a**).^62^ This database tracks biologically-relevant protein-ligand structures and binding interactions, and is updated on a weekly basis. In total, 854 complexes were successfully predicted and subsequently compared. When aligning structures based on pocket residues within 5Å of the ligand, AF3 correctly inferred the binding pocket in 98.2% of cases (pocket RMSD < 2Å). However, the success rate drops to 49.6% (424 out of 854 complexes) when requiring simultaneous success in both ligand pose (ligand RMSD < 2Å) and protein pocket reproduction (pocket RMSD < 2Å) (**Fig. 3a**, upper panel).

To test whether performance increases with higher 2D fingerprint similarity, we expanded our analysis to include all protein-ligand complexes deposited in the PDB during the same time window, without imposing the novelty filter. In total, 8,076 complexes were successfully predicted and successfully analyzed. A portion of complexes failed at either the prediction stage or analysis stage, and these entries were discarded. See **Methods** for a detailed breakdown. We then plotted ligand pose success rates (ligand RMSD < 2Å) as a function of Tc similarity to the AF3 training set (**Fig. 3a**, bottom panel). Though we expected a higher 2D similarity to correlate with improved pose accuracy, the success rate remains relatively stable, with only marginal gains at higher similarity levels (**Fig. 3a**, bottom panel). This indicates that AF3 success does not rely on memorizing topologically similar ligands. Instead, AF3 achieves similar success across a broad range of ligand similarity values.

Similar to the DUDE-Z benchmark, we next examined whether pose prediction accuracy correlates with ligand-only pTM. We hypothesized that ligand-only pTM could reflect the extent to which AF3 encodes or “memorizes” chemical features of the ligand independent of the protein. To test this, we ran AF3 for all 8,076 complexes using only the ligand SMILES as the input, omitting the protein sequence entirely.

Similar to the trend observed with topological similarity, ligand-only pTM showed no strong relationship with pose success rate (**Fig. 3b**, bottom panel, grey curve). A modest dependence emerged only when the ligand-only pTM metric was multiplied by the binding pocket similarity score (PS-Score) (**Fig. 3b**, bottom panel, purple curve). The PS-Score quantifies geometric and sequence similarity among pocket residues within 5 Å of the ligand.^54^ When comparing novel predictions (ligand-only pTM * PS-Score ≤ 0.35), 1,637 out of 4,152 complexes (39.4%) achieved both ligand RMSD and protein RMSD < 2 Å (**Fig. 3b**, upper panel). In contrast, 59.5% of predictions fell into the “correct pocket but incorrect ligand” category, indicating that ligand placement remains the primary failure mode. Notably, although ligand-only pTM performs well when distinguishing computationally generated ligands from decoys, it is far less informative for experimentally solved complexes. In these cases, ligand-only pTM confidence declines when the protein sequence is omitted, suggesting that AF3 does not encode ligand features in isolation but instead relies on protein context to reliably reconstruct ligand geometry and pose.

We also reproduced the memorization effects reported by Škrinjar et al.^51^, where the team identified that success rate is strongly correlated with 3D ligand similarity and pocket similarity. Because our benchmark set has a 30% overlap with the Runs N’ Poses dataset, we expected to observe similar behavior. Importantly, all complexes in our study were deposited after AF3’s validation cutoff, minimizing any influence from model-selection bias. To quantify these effects, we evaluated 3D ligand similarity and pocket similarity using the same conceptual metrics as prior work, but with slight methodological differences (see **Methods**). Using ComboROCS alone as the similarity metric revealed a modest memorization trend (**Fig. 3c**, bottom panel, grep curve), but is strengthened when multiplied by the PS-Score. This multiplied score produces a combined ligand-pocket 3D similarity metric. Among all similarity measures tested, this composite metric displayed the clearest memorization dependence (**Fig. 3c**, bottom panel, green curve and **Extended Data Fig. 8b**), suggesting that it is the best pre-run predictor of AF3 success. When applying the same novelty threshold based on this combined metric (≤ 0.35), only 173 out of 970 complexes (17.8%) achieved both ligand RMSD < 2 Å and pocket RMSD < 2 Å (**Fig. 3c**, upper panel). Most remaining cases (78.7%) had correctly predicted pockets but incorrect ligand placement. Therefore, for novel complexes jointly dissimilar in ligand shape/pharmacophore and pocket architecture, AF3 typically reconstructs the binding environment accurately but mispositions the ligand.

Although ligand RMSD < 2 Å is a widely used criterion for evaluating pose reproduction, this metric primarily measures geometric agreement and may not reflect whether biologically relevant interactions—such as hydrogen bonds, salt bridges, π–π stacking, and/or cation–π contacts—are preserved. Ligand RMSD is also sensitive to protein alignment, which can mask failures in interaction-level accuracy. As suggested by Errington et al.^63^, a more functionally relevant definition of pose accuracy is whether the native protein-ligand interaction network is recovered. To assess this, we used the interaction profiler LUNA^64^ to compute protein–ligand interaction fingerprints (PLIFs) and compared predicted complexes to their experimental counterparts. As shown in **Fig. 3d**, we plotted PLIF recovery versus ligand RMSD for all predicted complexes, using a PLIF recovery success threshold of 0.8 (i.e., ≥80% of native interactions are reproduced). Among the 4,235 complexes with ligand RMSD < 2 Å, 62% (2,622/4,235) failed to recover at least 80% the native interactions. Even when AF3 places the ligand near the correct location, binding interactions are not always reproduced. When applying this interaction-based criterion across different similarity measures (**Fig. 3e** and **Extended Data Fig. 8c**), neither high 2D nor 3D similarity corresponded to improved interaction recovery. Together with the observation that the strongest memorization effect arises from ligand–pocket similarity rather than interaction similarity, we conclude that AF3’s pose success derives largely from spatial memorization rather than learning transferable rules of binding. AF3 remembers where ligands go, but not why they bind.

### Prospective Screen of σ_2_ Receptor

Despite its decent performance on retrospective benchmarks, AF3’s capacity for prospective ligand discovery has not, to our knowledge, been tested, especially on a novel target outside its training set. To address this gap, we performed a head-to-head screening campaign using both the physics-based docking engine DOCK3 and the co-folding–based method AF3 to screen the Enamine in-stock library against the σ_2_ receptor. We selected the σ_2_ receptor for two reasons: i) its experimental structures were deposited to the PDB after AF3’s training cutoff date (09/30/2021), which serves as a trial for AF3 ligand discovery on novel targets, which mimics a novel discovery scenario in real-world drug discovery; and ii) in retrospective testing on the σ_2_ large experimental testing dataset, AF3 achieved an early enrichment better than DOCK3 and well reproduced ligand crystal poses bound to the σ_2_ receptor (**Extended Data Fig. 7**).

More than 690,000 monocations were sub-selected from the 4.6 million in-stock molecule library and docked against the σ_2_ receptor using both AF3 and DOCK3. The monocation filter was used because of computing resource limitations for AF3 inference. As a proof of concept, we evaluated enrichment by scoring the 133 known σ_2_ receptor actives from the previous large-scale experimental set against the full 690,000-compound monocationic prospective library. AF3 ipTM showed slightly better enrichment of known actives than AF3 mPAE (ipTM logAUC = 13.5, AUC = 74.1 vs. mPAE logAUC = 9.88, AUC = 69.2, **Extended Data Fig. 9**), so ipTM was selected as the ranking metric for the AF3 screen. Among the top 100,000 compounds from each campaign, DOCK3 yielded greater chemotype diversity, identifying 42,013 unique Bemis-Murcko scaffolds compared with 31,479 for AF3. We then applied consistent post-processing—novelty filtering to σ_2_ knowns, removal of compounds with unsatisfied hydrogen bond donors or acceptors, and diversity clustering—to the top 100,000 molecules from each screen, generating a prioritized list for purchase without any manual inspection (see **Methods** for details). In total, 108 top-ranking molecules from the DOCK3 screen and 114 from the AF3 screen were selected for purchase and testing (**Table S2**).

In primary radioligand displacement assays, 27 out of 108 molecules from the DOCK3 screen displaced over 50% of the [^3^H]-DTG-specific σ_2_ binding at 1 μM, a hit rate of 25%. In the secondary radioligand binding assays, the top 5 hits from the DOCK3 campaign had Ki values from 39 to 199 nM (**Fig. 4c**). Meanwhile, 15 of the 114 molecules from the AF3 screen met the same displacement threshold, resulting in a 13% hit rate, roughly half that of DOCK3. Across the 42 hits from both campaigns, each represented a unique Bemis–Murcko scaffold, with no scaffold shared between the AF3-and DOCK3-tested molecules, indicating that the two campaigns prioritize not only different molecules but also different families of molecules. Based on this lower hit rate, one might expect the top hits from the AF3 campaign to be weaker on average than those from DOCK3. Unexpectedly, the top five AF3 hits still exhibited comparable affinity distribution, with Ki values ranging from 13 to 131 nM (**Fig. 4c**), comparable to the 39 to 199 nM range observed for the top DOCK3 hits. Thus, although AF3 was less productive overall, the potency distribution among its best-scoring compounds remained competitive. These findings suggest that although AF3 can yield potent actives, its prospective performance on new targets remains limited compared to a physics-based approach. When combined with the cost and efficiency analysis, the differences are even more striking. The AF3 screen required 23,000 GPU hours on A100 GPUs, whereas DOCK3 completed the same task in 673 CPU hours on commodity hardware, a 34-fold difference in runtime compounded by a 200-fold difference in hardware price. Together, this translates to an overall efficiency gap of 6,700-fold in favor of DOCK3.

**Figure 4.**
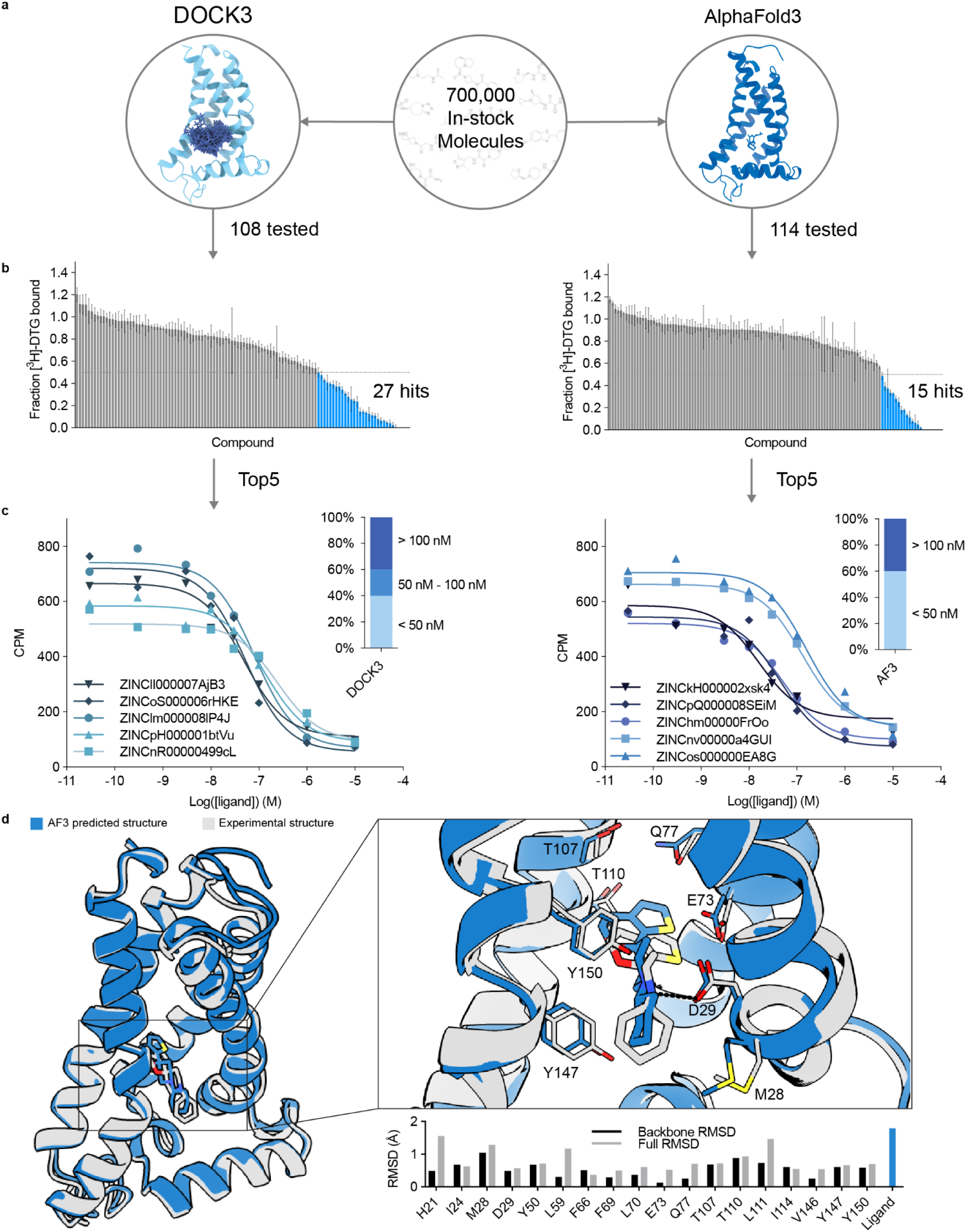
Prospective head-to-head σ_2_ receptor screening using AF3 and DOCK3. **a.** Schematic of the head-to-head virtual screening campaign. Over 700,000 monocationic compounds from the Enamine in-stock library were screened against the σ_2_ receptor using either the physics-based DOCK3 docking engine or the deep-learning co-folding model AF3. The top 100,000 ranked compounds from each screen were filtered for novelty and favorable interactions, then clustered by chemical diversity, yielding 108 DOCK3-selected and 114 AF3-selected molecules for experimental testing. **b.** Primary radioligand displacement results at 1 µM concentration. Blue bars indicate compounds that displaced >50% of [^3^H]-DTG binding, corresponding to experimentally confirmed hits. **c.** Secondary radioligand binding curves and Ki determinations for the top five hits from each campaign. Points shown as mean ± s.e.m. from three technical replicates. Inset bars summarize the affinity distribution of confirmed hits binned as < 50 nM, 50–100 nM, and > 100 nM. **d.** Structural validation of the AF3-predicted σ_2_ receptor complex with the AF3 top hit ZINCkH000002xsk4 against a newly solved X-ray crystal structure of the σ_2_ receptor bound to the same ligand. Blue, AF3-predicted receptor and ligand pose; gray, experimental crystal structure. Left, global superposition of the two complexes. Right, zoomed view of the orthosteric binding pocket, with key residues shown as sticks and the conserved salt bridge between the ligand’s protonated amine and D29 indicated by a dashed line. Bottom, per-residue root-mean-square deviation (RMSD, Å) between the AF3-predicted and experimental structures across binding-pocket residues, reported for backbone atoms (black) and all heavy atoms (gray); the rightmost pair of bars reports the ligand-pose RMSD between the AF3-docked and crystallographic poses of ZINCkH000002xsk4.

For all 222 newly tested compounds in the σ_2_ prospective screen, we first examined whether AF3’s ligand-only bias carried over to these newly tested σ_2_ compounds. As shown in **Extended Data Fig. 10a**, molecules with higher ligand-only pTM scores (>0.5) were in fact less likely to be active, defined here as > 50 % radioligand displacement, with hit rates dropping from 20 % to 13 %. This indicates that compounds with high complex-level confidence (ipTM) but also high ligand-only pTM are more likely to be false positives in experimental testing, suggesting that ligand-only pTM may be used as an additional filtering criterion in future prospective campaigns. We next assessed the physical plausibility of AF3- and DOCK3-generated complexes using PoseBusters. DOCK3 produced a slightly higher fraction of PoseBusters-valid predictions (107/108, 99 %) compared with AF3 (106/114, 93 %), and importantly, all PoseBusters-invalid compounds were experimentally inactive: 9 invalid inactives out of 180 total inactives (5% invalid rate), with 0 invalid actives. Thus, while PoseBusters flags only a small fraction of candidates, 100% of the flagged molecules are inactive in this dataset, indicating that structural implausibility can serve as a useful filter in prospective campaigns (**Extended Data Fig. 10b**). To further interrogate prospective enrichment beyond pose plausibility, we applied Boltz-2^33^, a recently released re-implementation of AlphaFold3 that extends co-folding with at least two additional features: a trained affinity prediction module that outputs (i) binding likelihood scores, and (ii) estimated log_10_(IC_50_) values. Benchmarking Boltz-2 against newly tested σ_2_ compounds from this study revealed that affinity predictions showed weak correlation with measured affinities and systematically overestimated potency for both AF3- and DOCK3-selected molecules (**Extended Data Fig. 10d**). Nonetheless, post-hoc rescoring with Boltz-2 improved enrichment, increasing the overall hit rate from 19 % (42 hits / 222 tested) to 30 % when filtering by binding likelihood cutoff of 0.8 (**Extended Data Fig. 10c**), and to 44 % when filtering by predicted affinity of log_10_(IC_50_) = -1 (**Extended Data Fig. 10d**). Thus, although Boltz-2 does not yet provide high-accuracy affinity predictions (**Extended Data Fig. 10e**), its binding classifier and affinity module still offer practical value as a downstream rescoring filter for prospective campaigns.

To directly evaluate the structural fidelity of the AF3 co-folding predictions that drove the prospective campaign, we determined the crystal structure of the σ_2_ receptor bound to ZINCkH000002xsk4, the most potent AF3-derived hit. We then overlaid this structure on the AF3-predicted complex for the same ligand (**Fig. 4d** and **Table 1**). Globally, the predicted and experimental receptor structures superpose closely, with a C_α_ RMSD of 0.67 Å. Within the orthosteric pocket, the side-chain rotamers of the key recognition residues are well reproduced. These include D29, which forms the conserved salt bridge with the protonated amine of ZINCkH000002xsk4, as well as the aromatic residues Y50, F66, Y147, and Y150 that line the cavity. Per-residue RMSDs are sub-Å for the backbone atoms of nearly all pocket residues and remain below 1.5 Å for full heavy-atom comparisons. The AF3-docked pose of ZINCkH000002xsk4 also closely matches its crystallographic binding mode, with a heavy-atom ligand RMSD of 1.8 Å. The pose recapitulates the salt bridge to D29 and the aromatic packing against Y147 and Y150 (**Extended Data Fig. 10f**). Together, these observations indicate that AF3 can produce both a near-native receptor pocket and a near-native ligand pose for a true active on a target outside its training set. The lower prospective hit rate relative to DOCK3 therefore appears to arise not from errors in the predicted complex of a true active. Rather, it reflects limitations in how the co-folding scoring function ranks and discriminates among candidate ligands once a high-quality pocket has been produced.

**Table 1.** Data collection and refinement statistics.

|  | **Bovine $\sigma_2$ receptor in complex with ZINCKH000002xsk4** |
| --- | --- |
| Wavelength | 0.4769 |
| Resolution range | 51.84 - 2.4 (2.75 - 2.4) |
| Space group | C 2 2 21 |
| Unit cell | 57.784 117.329 59.922 90 90 90 |
| Total reflections | 121470 (39154) |
| Unique reflections | 9101 (2959) |
| Multiplicity | 13.3 (13.2) |
| Completeness (%) | 99.59 (99.22) |
| Mean I/sigma(I) | 8.08 (0.71) |
| Wilson B-factor | 65.85 |
| R-merge | 0.2839 (5.795) |
| R-meas | 0.2958 (6.034) |
| R-pim | 0.08183 (1.667) |
| CC1/2 | 0.998 (0.551) |
| CC* | 0.999 (0.843) |
| Reflections used in refinement | 8237 (2672) |
| Reflections used for R-free | 406 (132) |
| R-work | 0.2367 (0.3761) |
| R-free | 0.2770 (0.3887) |
| Number of non-hydrogen atoms | 1452 |
| macromolecules | 1393 |
| ligands | 20 |
| solvent | 4 |
| Protein residues | 166 |
| RMS(bonds) | 0.009 |
| RMS(angles) | 1.08 |
| Ramachandran favored (%) | 96.34 |
| Ramachandran allowed (%) | 3.66 |
| Ramachandran outliers (%) | 0.00 |
| Rotamer outliers (%) | 1.32 |
| Clashscore | 7.88 |
| Average B-factor | 79.83 |
| macromolecules | 79.60 |
| ligands | 86.13 |
| solvent | 75.93 |

Statistics for the highest-resolution shell are shown in parentheses.

## Discussion

Deep-learning co-folding models such as AF3 have introduced the possibility that protein-ligand complexes can be predicted directly from sequence and ligand inputs, raising the question of whether such models might serve not only as structure predictors but as AI-based docking engines. Despite the excitement surrounding AF3, its practical application for ligand discovery has remained unclear. This study set out to evaluate AF3’s potential for structure-guided ligand discovery. Rather than just asking whether AF3 can accurately generate protein–ligand complexes, we focused on the question that matters for ligand discovery: under what conditions, and to what extent, can AF3 accomplish tasks in ligand pose prediction, enrichment of actives from a chemical library, and prospective hit identification. First, in retrospective screening on DUDE-Z, AF3 can reliably recover crystallographic ligand poses when supplied with protein templates, achieving <2 Å ligand RMSDs in 86% of targets. On the enrichment benchmark, AF3 appeared to outperform the physics-based docking program, DOCK3, across 43 drug targets, but this advantage was largely from hidden ligand-only biases in decoy generation rather than from genuine ligand-protein complex prediction. When the same comparison was repeated on three large experimental datasets (σ_2_, D_4_, AmpC) containing >2,500 tested compounds and free from such biases, DOCK3 achieved stronger overall enrichment, while AF3 contributed mainly to early enrichment. Second, when evaluated on >8,000 protein–ligand complexes deposited after AF3’s training and validation cutoff, AF3 pose accuracy declined sharply for ligands and pockets that were structurally novel, revealing strong dependence on training-set similarity and indicating that AF3 memorizes spatial binding poses rather than learning transferable rules of molecular recognition. This suggests that AF3 is likely to perform well only for complexes similar to those represented in its training set. The memorization curve also provides a practical model for users to estimate or set expectations for AF3’s performance on their own complexes. Finally, we compared AF3 with DOCK3 in the first prospective head-to-head virtual screen against the σ_2_ receptor, which is absent from the AF3 training set. Although the AF3 campaign required orders of magnitude more computational resources than docking, its hit rate was still two-fold lower (13% vs. 25%). However, despite the lower hit rate, AF3 was still able to identify a potent 13 nM binder, and the top-ranking AF3 hits showed comparable affinities to those obtained from physics-based docking. An X-ray crystal structure of the σ_2_ receptor bound to the top AF3-derived hit, ZINCkH000002xsk4, further showed that the AF3 prediction recovered both the ligand binding pocket and the ligand pose (heavy-atom ligand RMSD of 1.8 Å) for a true active on a target outside AF3’s training set, indicating that the lower prospective hit rate reflects limitations in scoring and ranking rather than in modeling the complex of an active. Together, our results reveal that AF3 performs well when the target and ligand chemotypes resemble its training set, but its generalization and scalability remain limited compared with traditional molecular docking. These observations define the current boundaries of the co-folding methods’ capabilities and frame where it might contribute most effectively within hybrid workflows rather than as a stand-alone replacement for docking. Our findings are reinforced by a contemporaneous study from Kim et al., which evaluated AF3, Chai-1, and Boltz-2 on 557 newly determined SARS-CoV-2 Mac1 complexes and re-analyzed the same σ_2_, D_4_, and AmpC datasets used here, reaching convergent conclusions from independent benchmarks^65^.

Historically, physics-based docking has shown substantial improvements in hit rate and affinity as library size grows^58–60,66,67^. Ultra-large docking of 490 million make-on-demand molecules against the σ_2_ receptor yielded a 51% hit rate and a best binder of 1.8 nM directly from docking^14^, and docking the same library against an AF2-predicted σ_2_ structure produced similar results (54% hit rate, 1.6 nM binder)^25^. In contrast, the much smaller in-stock library used here resulted in a 25% hit rate and 32 nM best binder for DOCK3, and a 13% hit rate and 13 nM best binder for AF3. Thus, AF3 achieved roughly half the hit rate of DOCK3 under identical screening conditions. More importantly, AF3 cannot yet access the library sizes that drive docking success: the GPU cost of AF3 inference makes >10^7^-scale campaigns infeasible at present. These results reinforce a central distinction: docking outcomes improve as ever larger and more stereochemically diverse chemical spaces are explored, whereas AF3’s current performance is limited by computational throughput. Future accelerations or generative-model-based narrowing of search space like Boltz-2 may partially alleviate this, but their impact remains to be demonstrated^50,68–70^.

A challenge for evaluating deep-learning models is that the boundary between “training” and “test” sets is inherently temporal: new protein–ligand complexes are deposited into the PDB weekly, and different co-folding models are trained with different cutoff dates. As a result, memorization cannot be evaluated once against a single fixed benchmark; the benchmark itself must evolve. To address this, we developed a workflow in this study for constructing post-training test sets and quantifying ligand-level similarity (via ComboROCS) and pocket-level similarity (via APoc). The full pipeline, including data curation scripts, is available as an open-source resource at https://github.com/lyulab/benchmarking-af3. We anticipate that such dynamically updating benchmarks will play a key role in separating true generalization from training-set recall as co-folding models continue to improve.

Several caveats merit mention. Our prospective σ₂ campaign focused on mono-cationic molecules, reflecting the dominant charge preference of previously reported σ₂ ligands. Although neutral binders exist (e.g., Cholesterol^14^ and 20S-hydroxycholesterol^71^), AF3 ranked almost exclusively cationic molecules: among the 506 σ₂ compounds tested, no neutral ligand appeared in the top 1% or top 5% of AF3 ipTM scores, and only 1% appeared in the top 10%, indicating a strong enrichment bias toward positively charged chemotypes. In addition, σ₂ was chosen because it is novel to AF3’s training set, but there are likely cases where AF3 would outperform docking—for example, targets with binding sites that bind large and flexible molecules, where the combinatorial explosion of conformational degrees of freedom poses challenges for sampling of physics-based approaches. However, such scenarios are difficult to test empirically, as most commercially accessible chemical libraries are designed under Lipinski’s “rule of five,”^72^ which inherently limits molecular size and flexibility, biasing available compounds toward small, drug-like molecules.

AF3 also has clear advantages in accessibility. Unlike docking, which often requires force-field parameterization, search-space tuning, and user expertise, AF3 can be run with minimal setup. However, the computational costs are substantial: co-folding methods’ training requires roughly 128 A100 (80GB) GPUs for one month^35^, which costs about $400,000 using cloud computing, and inference is slower and more resource-intensive than CPU-based docking. Under these cost constraints, one might expect co-folding methods to deliver clear performance gains over docking, yet our retrospective and prospective evaluations do not support that assumption. Instead, AF3 offers complementary strengths, ease of use, template-free complex generation, and early enrichment value, rather than across-the-board victories.

Looking ahead, several directions may help realize the full potential of deep-learning co-folding. First, expanding the diversity of protein–ligand complexes in training—especially those structurally dissimilar to current PDB entries—will likely yield greater improvements than architectural refinements alone. Two ongoing efforts point in this direction: [AI Structural Biology (AISB) Network](https://www.apheris.com/join-a-network/aisb), which enables secure federated learning on proprietary pharmaceutical structure data, and [OpenBind](https://www.diamond.ac.uk/Home/News/LatestNews/2025/Diamond-Light-Source-will-host-a-new-pioneering-AI-driven-drug-discovery-consortium.html), which aims to generate 500,000 new public protein–ligand structures in the next five years. One of the more recent co-folding methods, Pearl from Genesis Molecular AI^39^, even explored using synthetic data for training, leading to better pose reproduction results than AF3. Second, co-folding outputs may be most effective not as replacements for docking but as rescoring layers on top of docking. For example, screening ultra-large libraries with DOCK3 for scale, then apply secondary rescoring methods, such as AF3 complex confidence metrics (ipTM or mPAE), ipSAE (if possible for small molecules)^73^, or the Boltz-2 affinity module, re-rank the top 1–5%. Both our retrospective D_4_ and σ_2_ analyses and our prospective σ_2_ campaign support this hybrid strategy, which improved hit rates relative to docking alone. Finally, the most fundamental need is architectural: future co-folding models must learn molecular recognition, not just memorize spatial arrangements dependent on training similarity.

These caveats should not obscure the central observations from this study. These findings demonstrate the training set dependence of AF3 in its successful prediction of protein-ligand complexes, while its retrospective and prospective performances in our current study provide optimism that co-folding can advance future drug discovery campaigns.

## Extended Data Figures

**Extended Data Figure 1.**
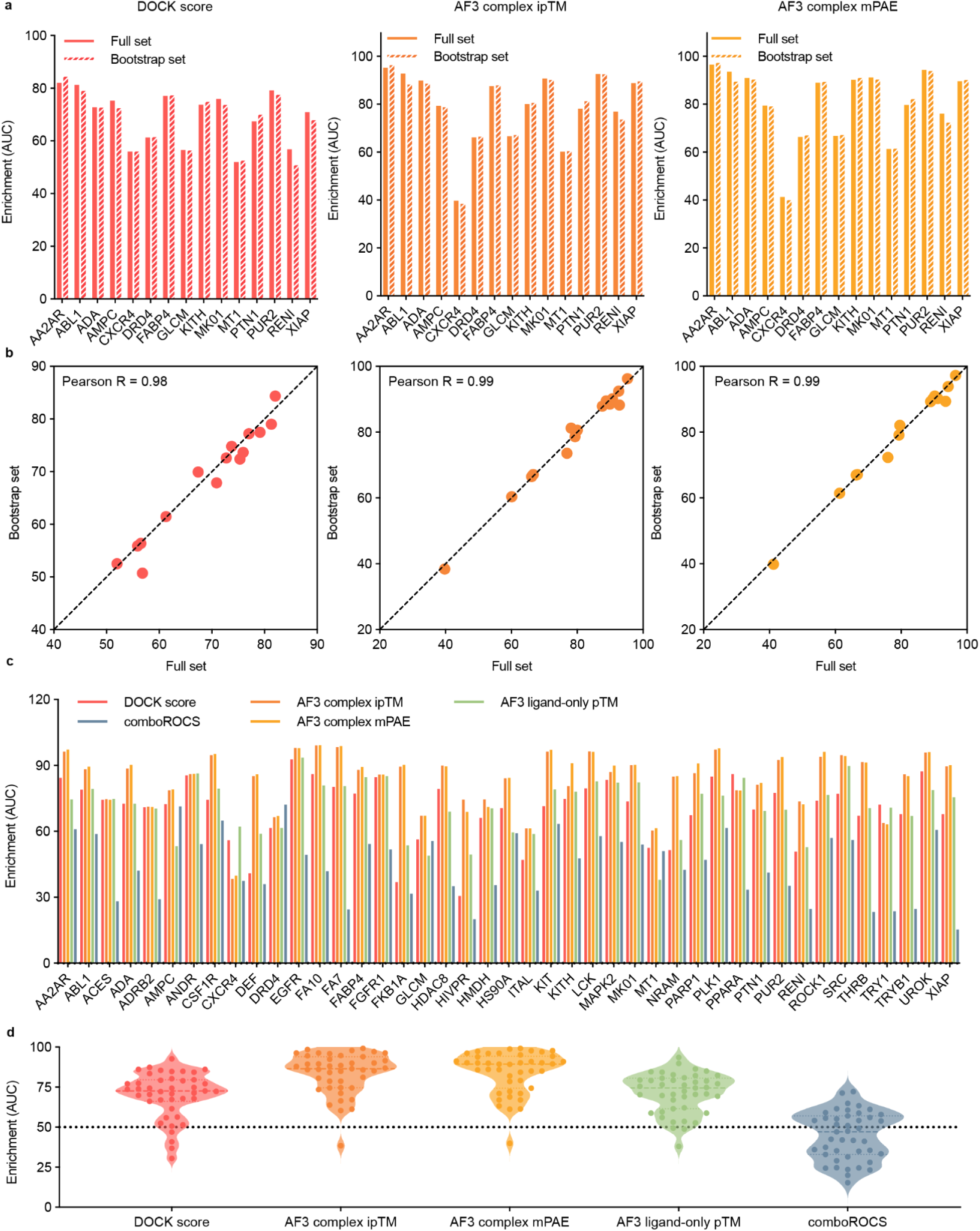
Validation and comparison of AF3 enrichment performance on the DUDE-Z benchmark (AUC evaluation). **a.** Enrichment of actives over decoys (AUC) for DOCK3 docking scores (left) and AF3 confidence metrics (ipTM, mPAE; middle and right) across 15 DUDE-Z targets. Solid bars indicate the full DUDE-Z dataset; hatched bars represent the down-sampled bootstrap subset (50 actives + 500 decoys per target). **b**. Correlation between enrichment (AUC) from the full and bootstrap datasets for DOCK3 (left), AF3 ipTM (center), and AF3 mPAE (right). Each point corresponds to one protein target. **c.** Comparison of AUC values across five scoring schemes—DOCK3 docking scores, ComboROCS 3D ligand similarity, AF3 complex metrics (ipTM, mPAE), and AF3 ligand-only pTM—across all 43 DUDE-Z targets. **d.** Distribution of target-wise enrichment (AUC) across five scoring schemes: DOCK3 docking scores, AF3 complex ipTM, AF3 complex mPAE, AF3 ligand-only pTM, and ComboROCS 3D ligand-shape similarity to AF3’s training set. Each point represents one DUDE-Z target; violin plots show the overall distribution across targets. The dashed horizontal line denotes the expected AUC for a random ranking.

**Extended Data Figure 2.**
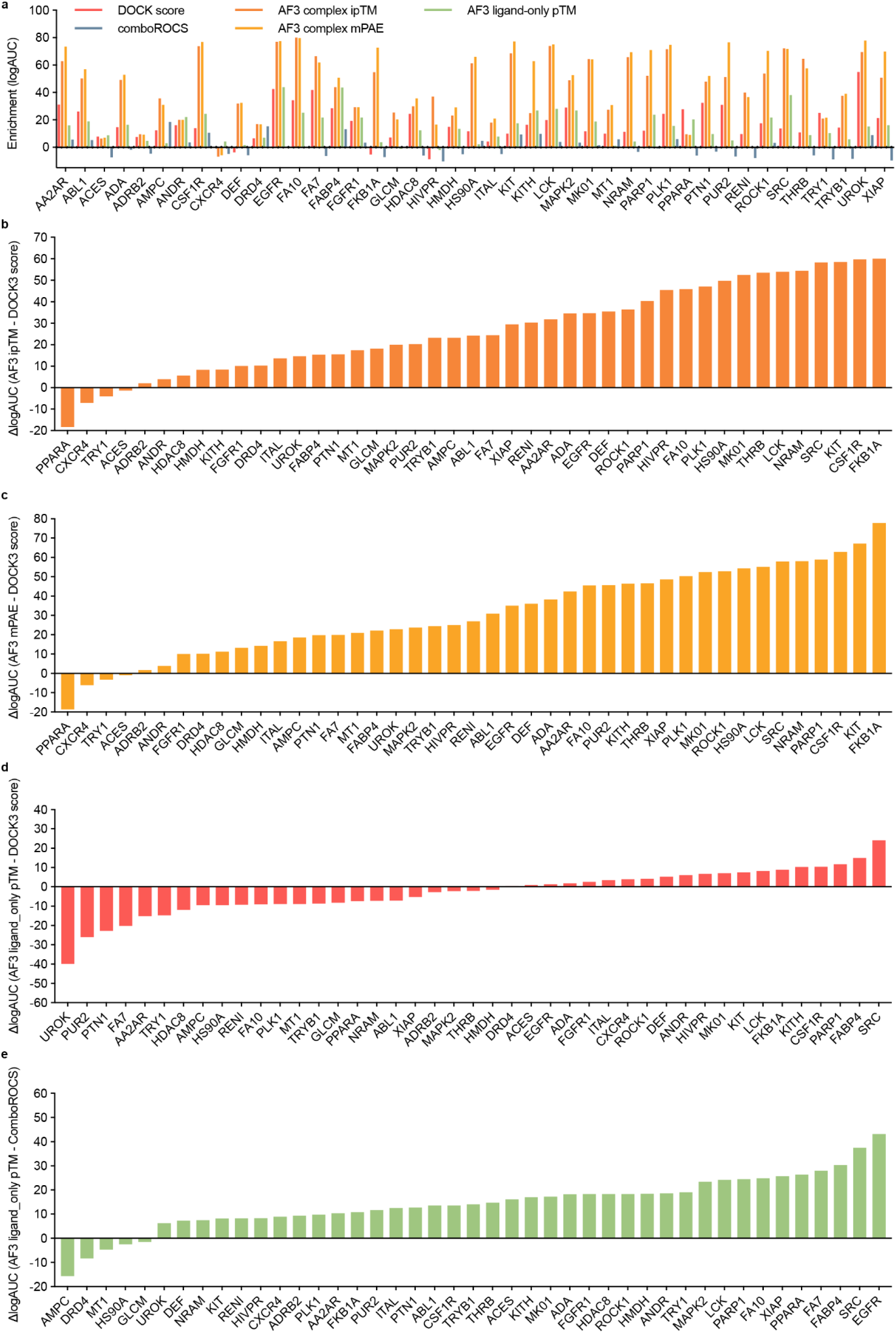
Comparison of enrichment (logAUC) differences between AF3 and DOCK3 across DUDE-Z targets. **a**. Target-wise enrichment comparison across four scoring schemes: DOCK3 docking scores, ComboROCS 3D ligand similarity, AF3 complex metrics (ipTM and mPAE), and AF3 ligand-only pTM. **b.** Difference in enrichment (ΔlogAUC) between AF3 complex ipTM and DOCK3 docking scores for each DUDE-Z target. Positive values indicate targets where AF3 ipTM achieved higher enrichment than docking. **c**. Difference in enrichment (ΔlogAUC) between AF3 complex mPAE and DOCK3 docking scores. **d.** ΔlogAUC between AF3 ligand-only pTM and DOCK3 docking scores. **e.** Difference in enrichment (ΔlogAUC) between AF3 ligand-only pTM and the 3D ligand similarity metric ComboROCS.

**Extended Data Figure 3.**
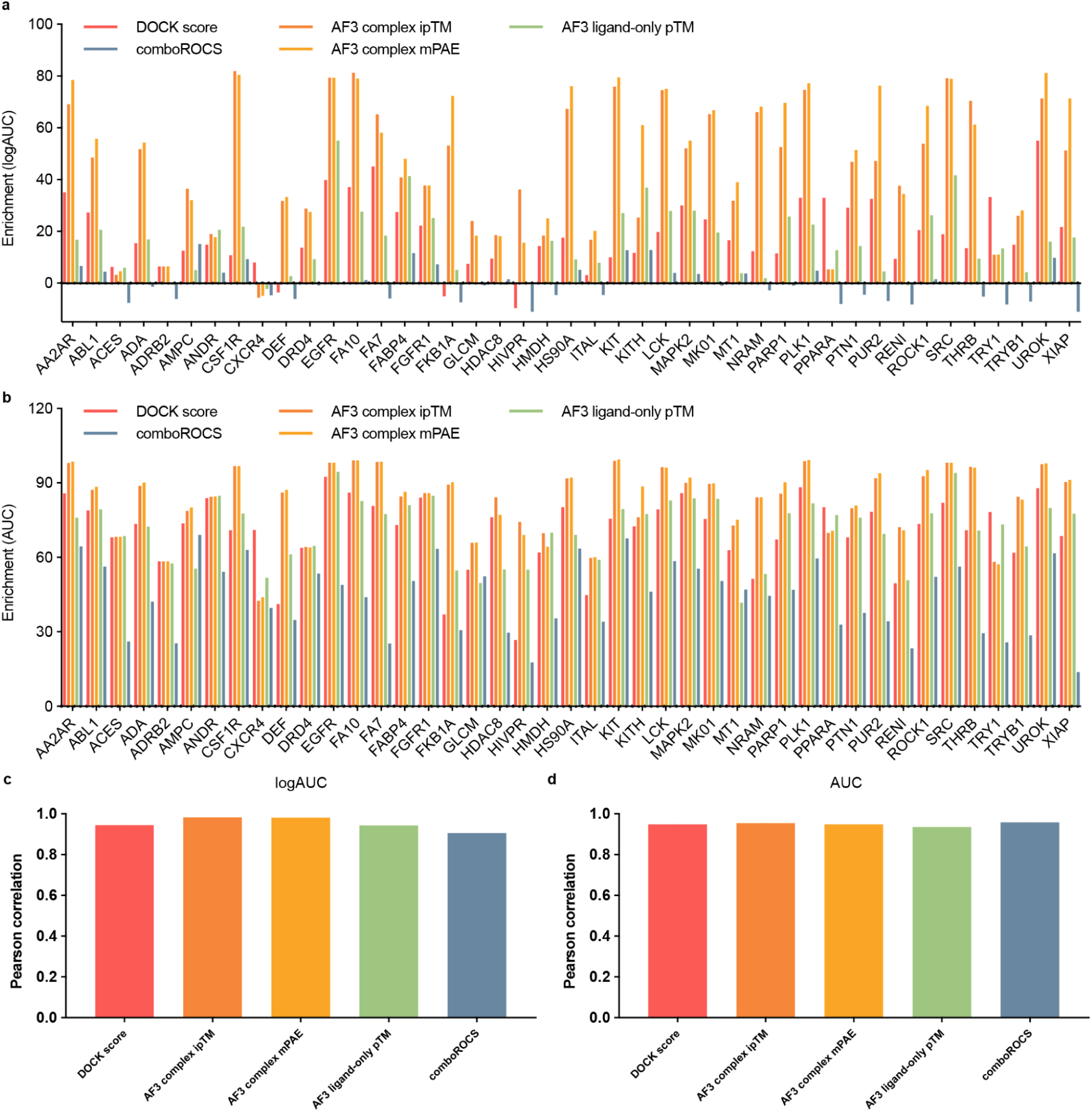
Enrichment performance of AF3 and DOCK3 on PoseBusters-validated molecules within DUDE-Z targets. **a**. Enrichment of actives over decoys (logAUC) for PoseBusters-validated ligand-protein complexes from each DUDE-Z target, comparing DOCK3 docking scores (red), ComboROCS ligand similarity (blue), AF3 complex ipTM (orange), AF3 complex mPAE (yellow), and AF3 ligand-only pTM (green). **b**. Corresponding enrichment (AUC) for the same set of PoseBusters-validated molecules evaluated across the same five scoring metrics. **c.** Correlation of logAUC values across the five scoring schemes (DOCK3 score, AF3 ipTM, AF3 mPAE, AF3 ligand-only pTM, and ComboROCS) from the full DUDE-Z ligand/decoy sets versus the PoseBusters-validated subsets. **d.** Corresponding correlation of AUC values.

**Extended Data Figure 4.**
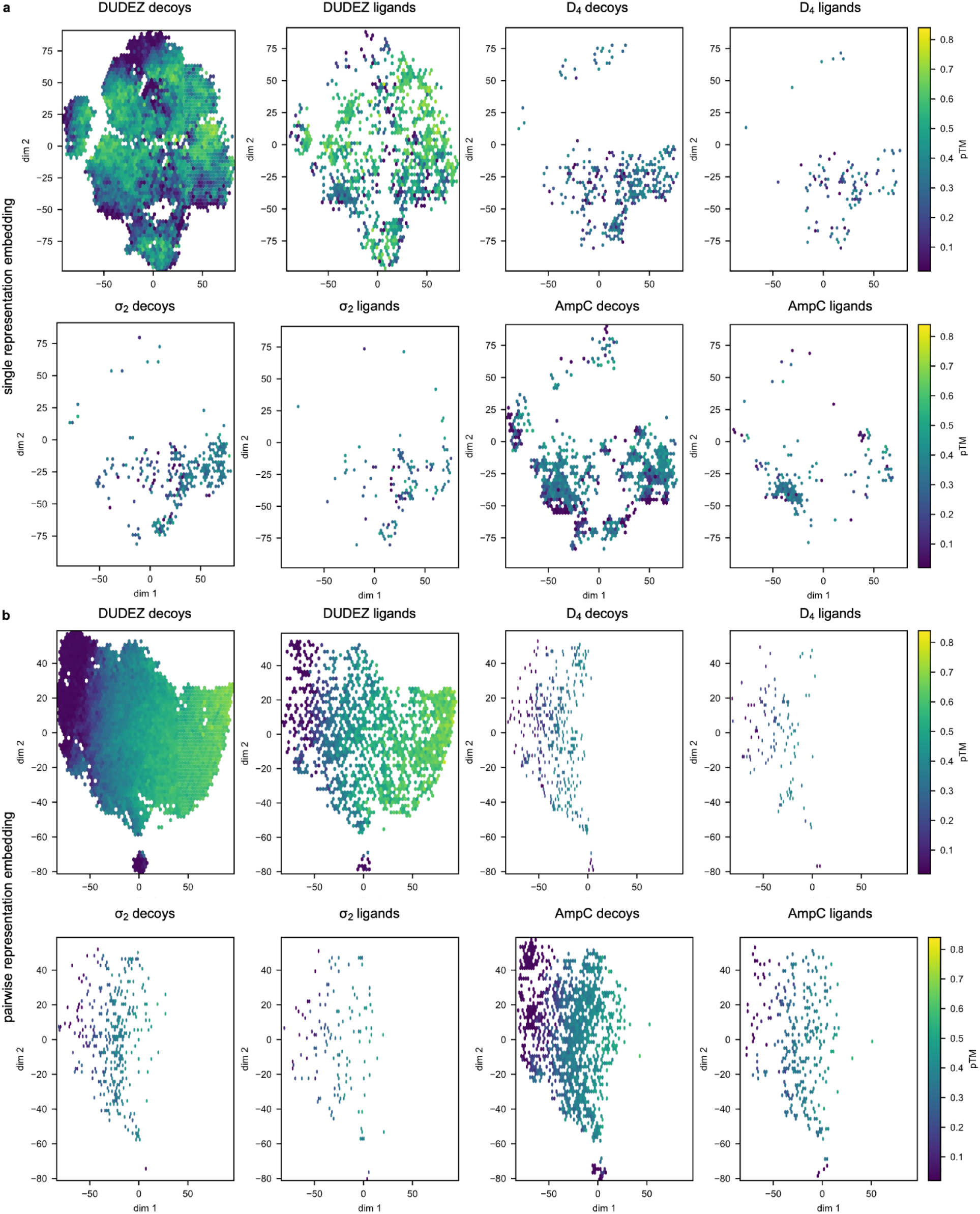
t-SNE projections of AF3 ligand-only single and pairwise representation embeddings colored by predicted pTM. **a.** Single-representation embeddings. For each compound in the DUDE-Z, D_4_, σ_2_, and AmpC datasets, AF3 ligand-only runs produce a single-representation embedding (256 tokens × 384 features). These embeddings were converted to fixed-length ligand fingerprints by mean-pooling across non-padding token embeddings. A single t-SNE projection was then learned on fingerprints from all compounds across all four datasets, and the resulting 2D embedding is plotted separately for each dataset. Within each dataset, decoys (left) and ligands (right) are shown as points colored by the AF3 ligand-only predicted pTM. **b.** Pairwise-representation embeddings. For the same compounds, AF3 ligand-only pairwise-representation tensors (256 tokens × 256 tokens × 128 features) were condensed to fixed-length fingerprints by mean-pooling all off-diagonal pairwise interaction features. As in a, a single t-SNE projection was fit using fingerprints from all four datasets, and the shared 2D embedding is visualized separately for each dataset, with decoys (left) and ligands (right) colored by ligand-only predicted pTM.

**Extended Data Figure 5.**
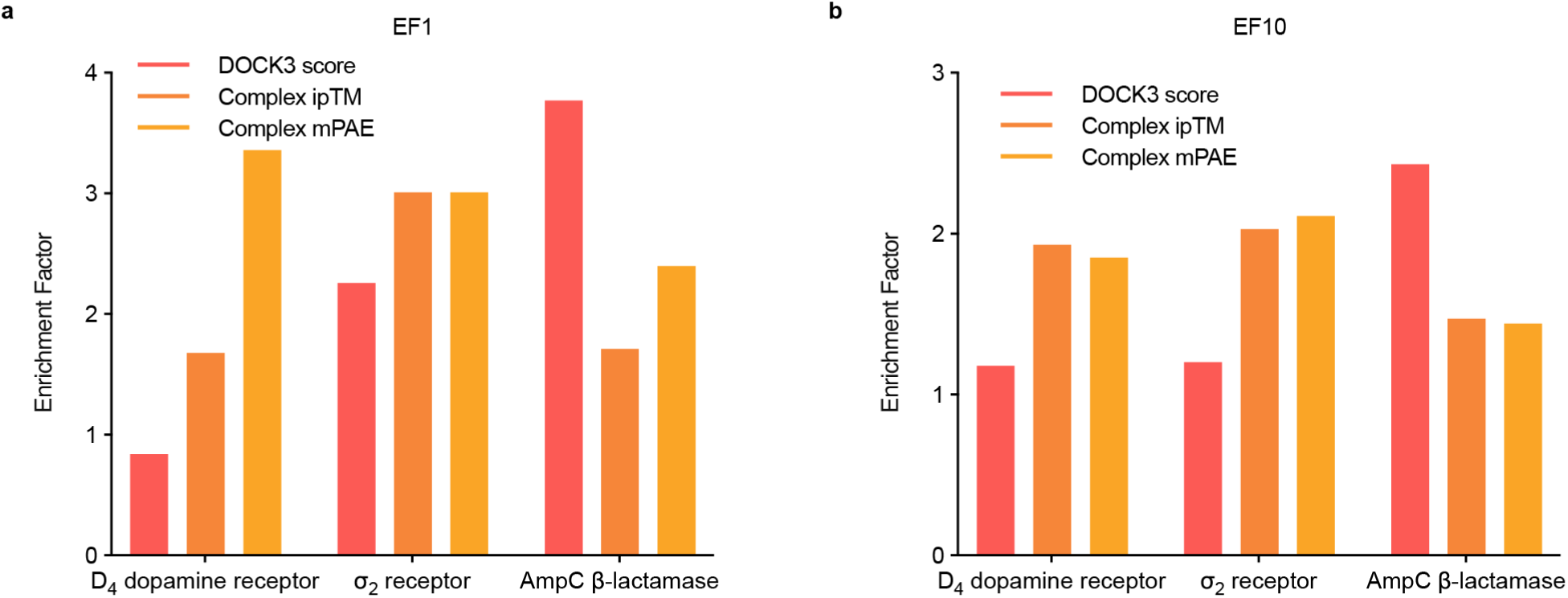
Enrichment factors of DOCK3 and AF3-derived complex metrics across three large experimental testing sets. **a.** Enrichment factor at the top 1% (EF1) and **b.** at the top 10% (EF10) of the ranked libraries for the D_4_ dopamine receptor, σ_2_ receptor, and AmpC β-lactamase campaigns. For each target, molecules were ranked by three scoring metrics: DOCK3 score (red), AF3 complex ipTM (pink), and AF3 complex mPAE (orange).

**Extended Data Figure 6.**
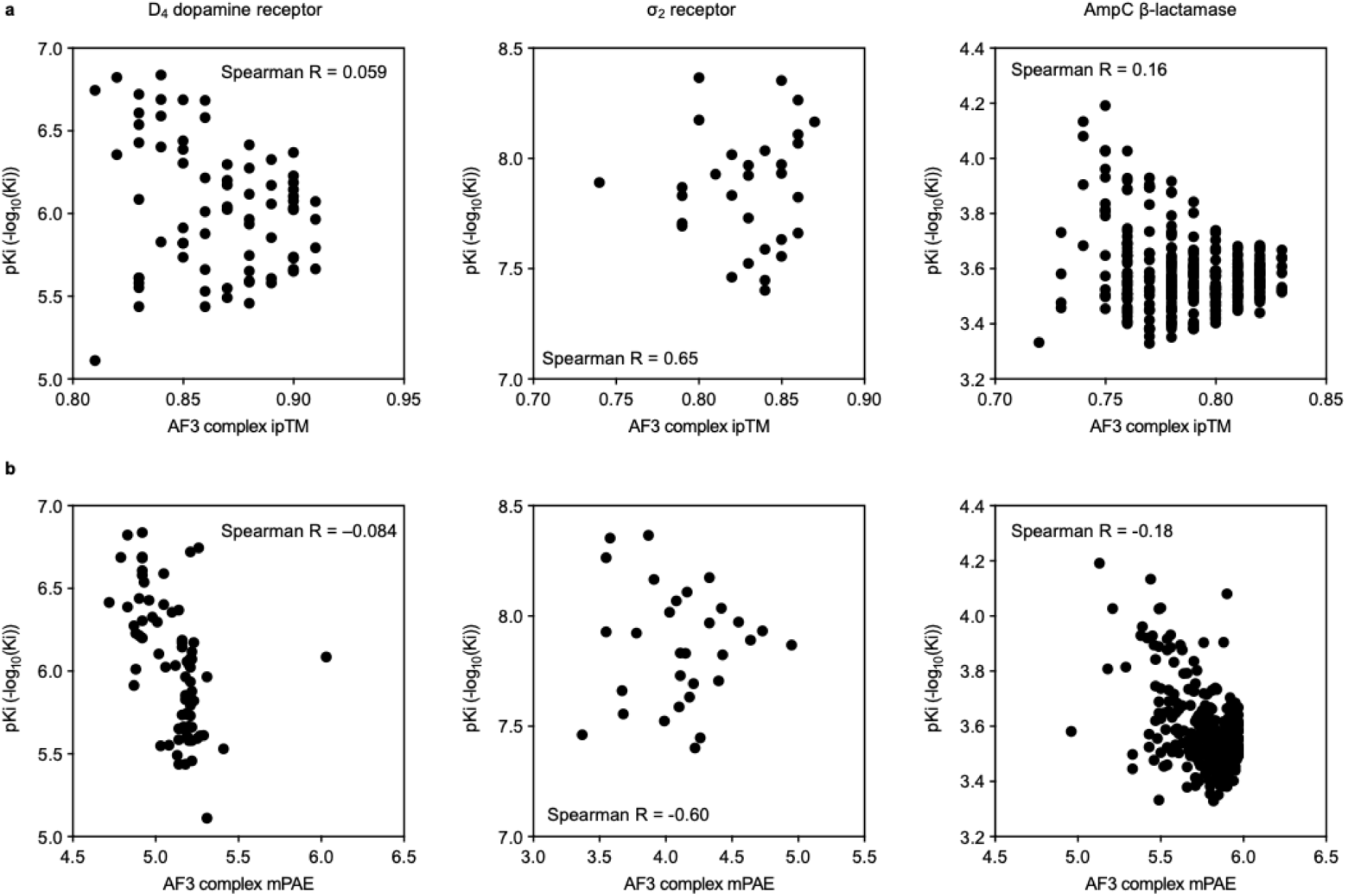
Correlation plots between AF3 confidence scores: **a.** AF3 complex ipTM; **b.** AF3 complex mPAE and experimental binding affinity (shown as pKi = -log_10_(Ki)) for three benchmark targets: D_4_ dopamine receptor (left), σ_2_ receptor (middle), and AmpC β-lactamase (right).

**Extended Data Figure 7.**
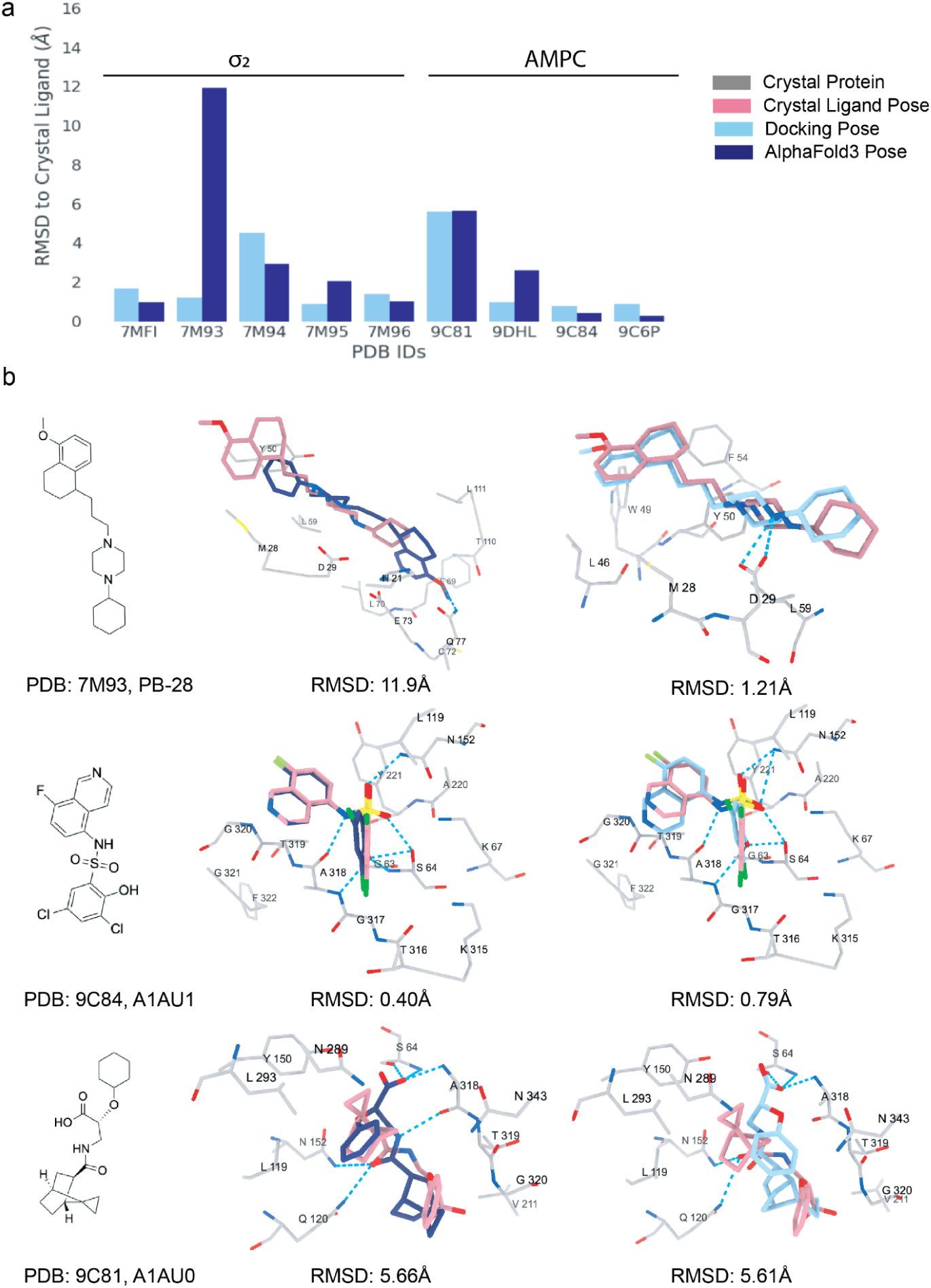
Comparison of AF3 and DOCK3 pose reproduction for post-training PDB complexes of σ_2_ and AmpC. **a.** Ligand RMSDs of AF3 (dark blue) and DOCK3 (light blue) predicted poses relative to crystallographic ligand poses for σ_2_ receptor and AmpC β-lactamase complexes deposited after the AF3 training cutoff (September 2021). **b.** Representative examples of ligand pose reproduction. Top: σ₂ receptor bound to PB-28 (PDB ID: 7M93), where the AF3 pose rotated the ligand 180° relative to the crystallographic pose (RMSD = 11.9 Å), while docking correctly reproduced the native orientation (RMSD = 1.21 Å). Middle: AmpC complex with A1AU1 (PDB ID: 9C84), where both AF3 and DOCK3 achieved near-perfect overlay (RMSDs < 1Å). Bottom: AmpC complex with A1AU0 (PDB ID: 9C81), where both AF3 and DOCK3 reversed the ligand orientation (RMSD = 5.66 Å and 5.61 Å, respectively) but maintained the same hydrogen-bonding network with residues S64, A318, Q120, and N152.

**Extended Data Figure 8.**
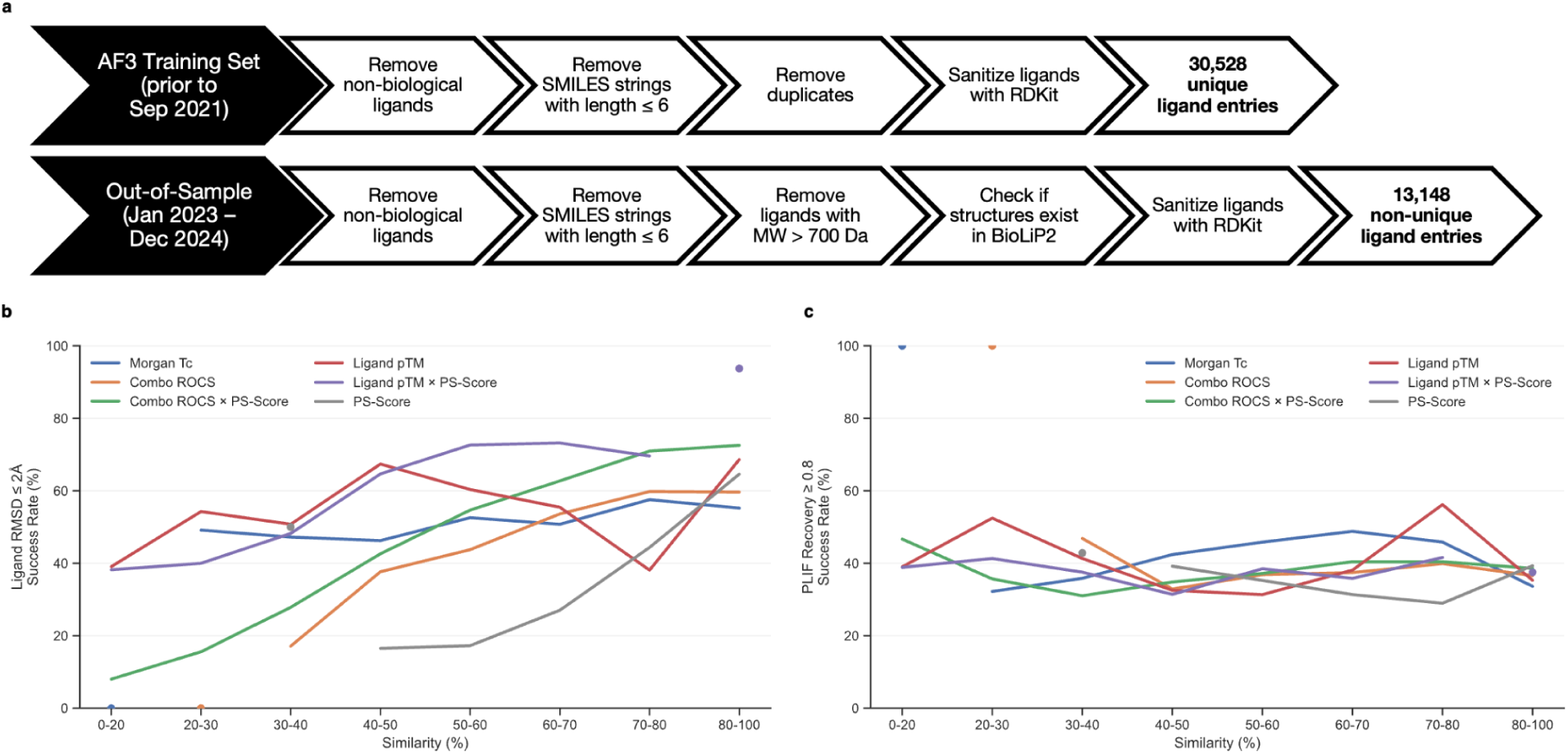
Success rates of AF3 ligand pose and interaction recovery as a function of similarity to the training set. **a.** Workflow used to create the out-of-sample dataset with data compilation and construction steps. Similarity metrics and novelty scores were later performed by comparing the out-of-sample set to the AF3 training set (details see **Methods**). **b.** Success rate of accurate ligand pose prediction (ligand RMSD ≤ 2 Å) across similarity bins defined by six metrics: 2D ligand similarity (Morgan Tc, blue), 3D ligand similarity (ComboROCS, orange), combined 3D and pocket similarity (ComboROCS × PS-Score, green), ligand-only pTM (red), ligand-only pTM × PS-Score (purple), and PS-Score alone (gray). **c.** Success rate of protein–ligand interaction recovery (Tversky index of interaction fingerprint ≥ 0.8) across the same similarity bins and metrics. Bins with fewer than 50 observations (n < 50) are displayed as single points without error estimates and line trends, as the sample size is insufficient to reliably calculate uncertainty.

**Extended Data Figure 9.**
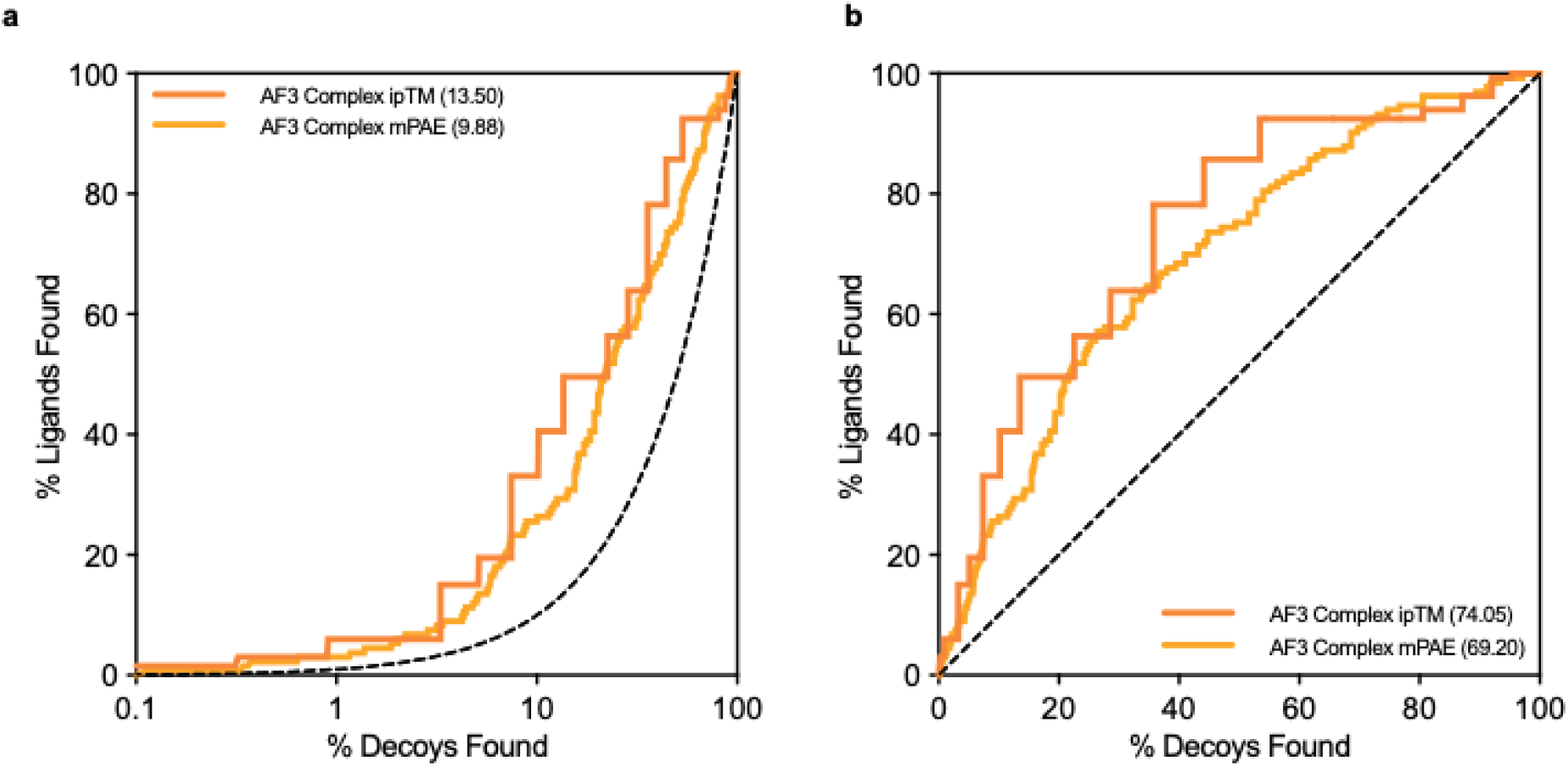
Enrichment of known σ₂ actives across the full monocationic library using AF3 confidence metrics. **a.** Log-adjusted ROC plot comparing enrichment of the 133 previously tested σ_2_ actives against the full 690,000-compound library using AF3 complex ipTM (dark orange) and AF3 complex mPAE (light orange). **b.** ROC plot for the same comparison.

**Extended Data Figure 10.**
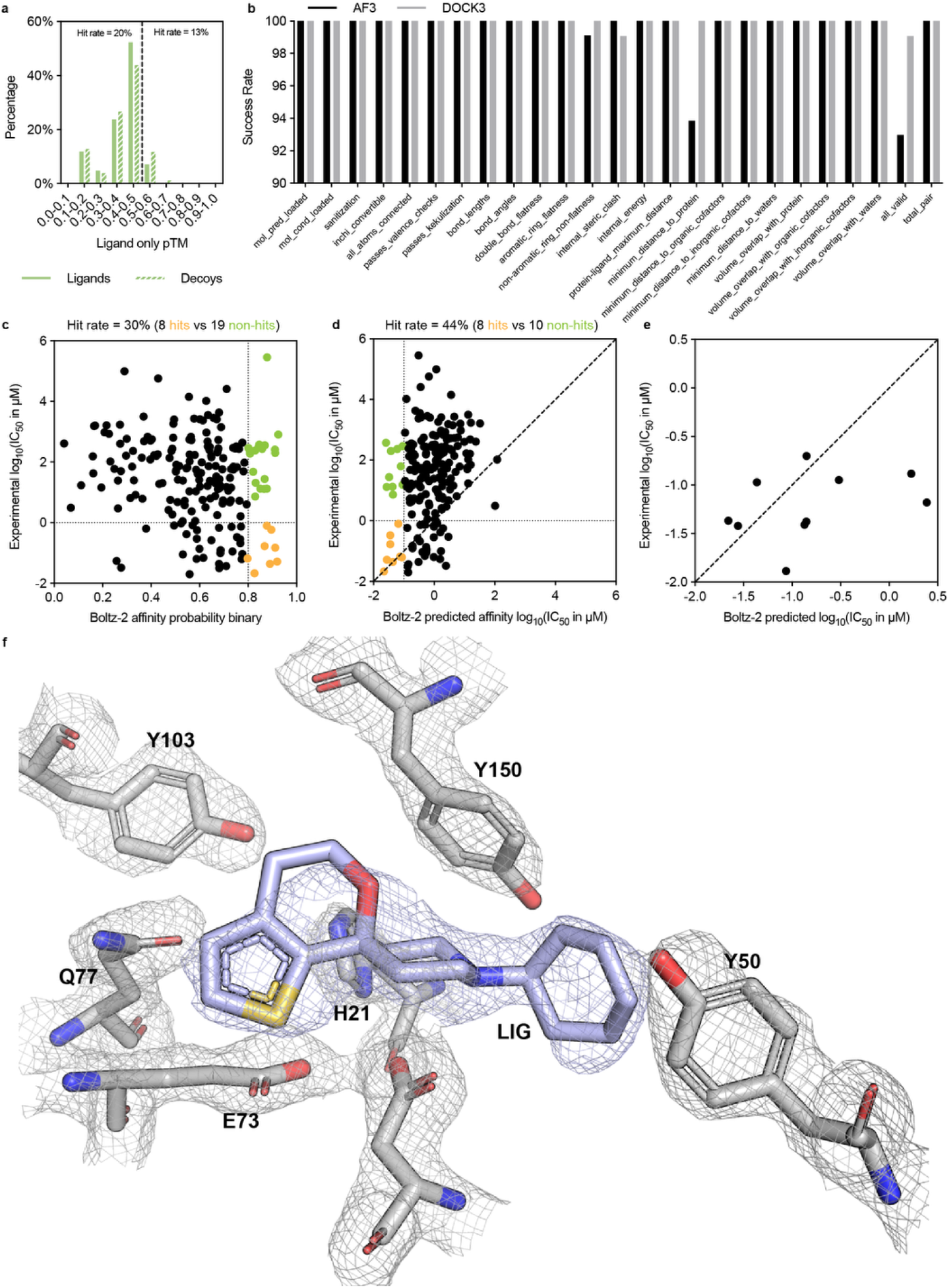
Ligand-only bias, PoseBusters validation, and Boltz-2 rescoring of prospective σ_2_ screening results. **a.** Distribution of ligand-only AF3 pTM scores for all 222 compounds tested in the σ_2_ receptor prospective screen. Molecules with higher ligand-only pTM (> 0.5) displayed a lower hit rate (13%) compared with those below the threshold (20%). **b.** PoseBusters validation success rates across all tested molecules from the DOCK3 and AF3 prospective campaigns. **c.** Experimental versus Boltz-2 predicted activity using the binary affinity probability metric.With an affinity probability cutoff of 0.8, Boltz-2 prioritized 8 active compounds among 27 tested, achieving a 30% hit rate. **d.** Comparison of experimental and Boltz-2 predicted log_10_(IC_50_) values using its affinity module. The model achieved a 44% hit rate (8 hits vs 10 non-hits). **e**. Correlation between Boltz-2 predicted and experimental affinities for the top σ_2_ hits with a dose response curve measurement. **f**. Crystal structure of the most potent AF3-derived σ_2_ hit (ZINCkH000002xsk4) bound in the orthosteric pocket, with the 2Fo–Fc electron density shown as a gray mesh. The ligand is engaged by surrounding binding-site residues, including H21, E73, Q77, Y50, Y103, and Y150.

## Methods

AlphaFold3 (version 3.0.0) model parameters were obtained with permission from Google DeepMind and inferences were conducted on The Rockefeller University high-performance computing cluster and the Delta system at the National Center for Supercomputing Applications. Multi-sequence alignment (MSA) was performed with the Jackhammer module from HMMER (version 3.4) if no templates were supplied. The example JSON input files used in this study are shown in SI. Five structures were sampled for each of five model seeds (4, 8, 9, 20, 31), and the best-performing structure by model ranking_score was used for analysis. For the ligand only AF3 prediction, the JSON input file only includes ligand smiles without any protein sequence information. DOCK3 ligand poses, docking scores, and experimental affinities were obtained from prior studies for AMPC (n=1,520), σ_2_ (n=506), and D_4_ (n=553). Additional (n=127) docking scores and activities for possible AMPC artifacts that were pre-filtered from the prior (n=1,520) study^74^ were obtained and analyzed with the larger AMPC test set.

### Evaluation of AF3 and DOCK3 on DUDE-Z

AF3 was run on all 43 DUDE-Z protein targets^53^. Templates were created from the corresponding PDB structures (corresponding PDB IDs available: https://dude.docking.org/targets) and PDB amino acid sequence data to bypass MSA. Enrichment of ligands over decoys was calculated using the enrich.py script from the DOCK3 source code^75^. Ligands in the AF3 training dataset, filtered for non-biological ligands (buffers and other solvents), proteolysis-targeting chimeras (PROTACs), and ions (cutoff: September 30, 2021, n=17,156 ligands) were downloaded, and their similarity to the AF3 training set was computed using OpenEye ROCS^57^. Ligand similarity is defined as the maximum ROCS similarity between the AF3 co-folded ligand pose and all ligands in the curated AF3 training set, reported as the sum of the Tanimoto coefficients for Color (pharmacophore features) and Shape (3D molecular volume), with values ranging from 0 to 2.

DOCK3.8 was re-run on all 43 DUDE-Z targets using their respective docking grid files (https://dudez.docking.org/DOCKING_GRIDS_AND_POSES) and precomputated ligand/decoy conformations (https://dudez.docking.org/Charge_matched_DUDE). Additionally, the INDOCK match_goal was set to 1000 to reduce docking runtime; the enrichment of ligands over decoys was calculated using the enrich.py script from the DOCK3 source code.

### Evaluation of AF3 against Large-Scale Experimental Testing Sets

Each ligand from the DOCK3 test set was co-folded with AF3 and evaluated with model confidence metrics: inter-chain predicted template modeling score (ipTM), a model-inferred value for template modeling TM score, and ranking score. Co-folded structures were aligned using APoc^54^ (version 1.0) with corresponding docking reference crystal structures using residues within 5 Å of any ligand heavy atom. The computed rotation and translation matrices from binding pocket alignment were applied to the co-folded ligands and root-mean-square-deviation (RMSD) to docked poses was calculated with the open source tool DockRMSD (version 1.1).^76^ Assayed activities (*K_i_* and displacement percentage) from previous virtual screening campaigns were used to calculate and compare enrichment between co-folding and docking. The *K_i_* cutoffs per compound were defined as in their original campaigns: 400 μM for AMPC, displacement greater than 50% [^3^H]-DTG for σ_2_, displacement greater than 50% [^3^H]-N-methylspiperone for D_4_. As a part of prior docking campaigns^14,60^, new crystal structures for σ_2_ (PDB: 7M93, 7M94, 7M95, 7M96) and AMPC (9C81, 9C84, 9DHL, 9C6P) were solved and used to compare co-folded and docked poses. The same alignment and RMSD software packages were employed in this analysis.

### Evaluation of AF3 on Out-of-Training-Set Targets

All structures released on or after January 13, 2023 were downloaded from the RCSB PDB (https://www.rcsb.org).^77^ Ions and very small fragments (ligand SMILES length 6 or less), very large ligands (more than a molecular weight of 700 Da), and other non-biological ligands were excluded, along with any structures not available in the BioLip2 database^62^ or unable to be processed with RDKit (https://www.rdkit.org). As stated in the RoseTTAFold All-Atom paper^78^ by Krishna et al., we also state non-biological ligands as a non-polymer entity that represent solvent or crystallization additives instead of true, biologically relevant binding partners. See **Supplementary Information** for more detail.

To quantify chemical and structural novelty relative to the AF3 training set, Tanimoto coefficients for ligand bitvectors (Morgan fingerprint radius of 2) and OpenEye similarity metrics were calculated between this filtered dataset and the AF3 training set, defined to be any structure released before September 30, 2021. To reproduce the memorization effects reported by Škrinjar et al., we computed comparable ligand-pocket 3D similarity metrics, but with two changes: (i) instead of using PLINDER’s Combined Overlap Score (SuCOS)^79^, we used OpenEye’s Rapid Overlay of Chemical Structures (ROCS)^57^ to calculate a Tanimoto Combo score (arithmetic sum of shape and color ROCS) termed “ComboROCS”; and (ii) instead of PLINDER’s binding pocket coverage score (“qcov_pocket”)^79^, we utilize the APoc package again to obtain the PS-Score. The obtained ComboROCS scores were normalized to [0,1].

Co-folded structures were aligned with their reference crystal structures using binding pocket residues within 5Å of any ligand heavy atom. Pocket similarity between inferred protein structures and experimental protein structures was calculated using the APoc software, which uses backbone geometry, side-chain orientation, and chemical similarity to construct a Pocket Similarity score (PS-score).^54^ The PS-score of each AF3-predicted structure was compared against the entire AF3 training set, and the maximum score was kept.

Novelty was defined strictly: a protein-ligand complex was considered novel if any of the above similarity metrics (Morgan Tc, comboROCS, PS-Score) fell below 0.35. Success in protein structure inference was defined as pocket residue RMSD to the protein crystal structure less than 2Å and success in ligand pose prediction was defined as crystal pose reproduction within 2Å. Success of interaction recovery was set at the 80% threshold (interaction fingerprint Tversky index ≥ 0.8). The LUNA toolkit^64^ was used to create an Extended Interaction FingerPrint (EIFP)^80^, and interaction fingerprint similarity between the experimental and AF3-predicted co-folded complexes was assessed using a one-sided Tversky index, weighted to measure the overlap relative to the reference fingerprint. Inspired by Errington et al.’s study on interaction recovery, we configured the LUNA workflow (https://luna-toolkit.readthedocs.io) to only look at interactions that are generally considered more biologically relevant: hydrogen and halogen bonds (donor and acceptor), π–π stacking (face-to-face and edge-to-edge), cation-π, and ionic interactions (anionic and cationic).^63^ All structures that have been successfully processed, predicted, and analyzed without hitting any failure points have been reported (n=8,076).

### Prospective Screens on the σ_2_ Receptor

Screening compounds in the Enamine in-stock library were pre-filtered for monocations, yielding 692,689 structures. The docking setup for σ_2_ was previously described, using a maximum strain energy per torsion angle of less than three units and total torsion strain less than eight units.^14,25,58^ AF3 co-folded structures were inferred as above, modified with a template based on the 7MFI PDB structure to bypass the multi-sequence alignment search. The top-ranking 100,000 compounds from each screen (sorted by energy for docking and ipTM for AF3) were filtered for novelty using the Morgan-based Tc against 2,232 σ_1/2_ ligands in ChEMBL (https://www.ebi.ac.uk/chembl) and 574 σ_2_ ligands from S2RSLDB (http://www.researchdsf.unict.it/S2RSLDB). Molecules with Tc ≥ 0.36 were eliminated, leaving 90,218 for DOCK3 and 76,251 for AF3. The remaining molecules were filtered for a salt bridge interaction with D29, three or fewer unsatisfied hydrogen bond acceptors, and zero unsatisfied hydrogen bond donors. The AF3 compounds were additionally filtered for total torsion strain and maximum strain per torsion angle as per the docking experiment. This resulted in 18,062 and 21,324 remaining compounds for docking and AF3 respectively. The interaction filter was implemented based on LUNA (https://github.com/keiserlab/LUNA) version 0.13.1. The remaining molecules were clustered with the Butina algorithm (RDKit ML Cluster, RDKit version 2024.03.5) using a distance cutoff of 0.68, resulting in 1,767 and 2,134 clusters for docking and AF3 respectively. The 120 top-scoring cluster members (by ipTM or docking energy) were prioritized for lab testing. Compounds with purity less than 90% were excluded, leading to 108 and 114 compounds successfully synthesized and ordered for lab testing. Other calculations, including Bemis-Murcko molecular scaffolds, were calculated with RDKit 2025.03.3.

### Membrane Preparation for Radioligand Binding Assay

Membranes for radioligand binding assays were prepared from Expi293 cells (Thermo Fisher Scientific) transiently transfected with wild-type σ_2_ receptor using FectoPRO (Polyplus-transfection) according to the manufacturer’s instructions. Cells were harvested 72 hours post-transfection and pelleted by centrifugation. Cell pellets were resuspended in 20 mM HEPES (pH 7.5), 2 mM magnesium chloride, and 1:100,000 (vol:vol) benzonase nuclease (Sigma Aldrich), supplemented with cOmplete Mini EDTA-free Protease Inhibitor Mixture Tablets (Roche). Cells were disrupted by Dounce homogenization and centrifuged at 50,000 × g for 20 min at 4°C. The resulting membrane pellet was resuspended in 50 mM Tris (pH 8.0), subjected to final Dounce homogenization, aliquoted, flash frozen, and stored at −80°C until use.

### Radioligand Binding Assays on the σ_2_ Receptor

For radioligand binding experiments membranes were thawed, homogenized, and incubated with 5 nM [^3^H]-DTG (PerkinElmer) in 100 μL binding reactions containing 50 mM Tris (pH 8.0), 0.1% (w/v) bovine serum albumin, and 50 nM PD-144418 to block σ_1_ receptor binding sites. For competition assays, competing ligands were added at 1 μM for single-point measurements or at indicated concentrations for full displacement curves. Reactions were incubated at 37°C for 2 hours with gentle agitation. Reactions were terminated by rapid filtration through glass fiber filters pre-treated with 0.3% polyethylenimine using a Brandel cell harvester. Filters were washed with ice-cold water, then individually soaked in 5 mL Cytoscint scintillation fluid (MP Biomedicals) overnight. Radioactivity was quantified using a Beckman Coulter LS6500 scintillation counter. All reactions were performed in triplicate using 96-well format. Data were analyzed using GraphPad Prism software (Version 10.6.1). K_i_ values were calculated by direct nonlinear regression fitting, with Cheng-Prusoff correction applied implicitly by the software. Estimated IC_50_ values were calculated using the equation IC_50_ = (1-%displacement) x [L]/(%displacement), where [L] = 1 μM based on single-point measurements.

### Protein expression and purification for crystallography

Bovine σ_2_ receptor was expressed and purified as previously described^14^. Briefly, bovine σ_2_receptor bearing an N-terminal protein C epitope tag followed by a 3C protease cleavage and truncated after residue 168 was expressed in Sf9 insect cells (Expression Systems) using the BestBac baculovirus system (Expression Systems) following the manufacturer’s protocol. Cells were infected at a density of 3× 10^6^ cells ml^-1^ and incubated with shaking at 27 °C for 60 h. Cells were then harvested by centrifugation, and the resulting pellets were stored at −80 °C until purification.

The ligand (ZINCkH000002xsk4) was included at 1 μM in all buffers throughout purification. Frozen cell paste was thawed and lysed by osmotic shock in 20 mM HEPES pH 7.5, 2 mM MgCl_2_, 1:100,000 (v:v) benzonase nuclease (Sigma Aldrich), and cOmplete EDTA-free Protease Inhibitor Cocktail (Roche). The lysate was centrifuged at 50,000 x g for 15 min. Pellets were resuspended and homogenized using a glass Dounce tissue homogenizer in 20 mM HEPES pH 7.5, 250 mM NaCl, 10% (v/v) glycerol, 1% (w/v) lauryl maltose neopentyl glycol (LMNG; Anatrace), and 0.1% (w/v) cholesterol hemisuccinate (CHS; Steraloids). Solubilization was carried out with stirring for 2h at 4°C. Insoluble material was removed by centrifugation at 50,000 x g for 30 min, and the clarified extract was supplemented with 2 mM CaCl_2_ before filtration through a glass microfibre filter (VWR).

The filtered solubilized membrane fraction was applied by gravity flow to 5 ml anti-protein C antibody affinity resin preequilibrated with 10 column volumes of 20 mM HEPES pH 7.5, 250 mM NaCl, 2 mM CaCl_2_, 1% (v/v) glycerol, 0.1% (w/v) LMNG, and 0.01% (w/v) CHS. A second wash was performed with 10 column volumes of 20 mM HEPES pH 7.5, 250 mM NaCl, 2 mM CaCl_2_, 0.1% (v/v) glycerol, 0.01% (w/v) LMNG, and 0.001% (w/v) CHS. Bound receptor was eluted with 20 mM HEPES pH 7.5, 250 mM NaCl, 5 mM EDTA, 0.1% (v/v) glycerol, 0.01% (w/v) LMNG, 0.001% (w/v) CHS, and 0.2 mg/ml protein C peptide.

Peak elution fractions were pooled, supplemented with 3C protease at a 1:100 (w:w) protease:receptor ratio, and dialyzed overnight at 4°C 20 mM HEPES pH 7.5, 250 mM NaCl, 2 mM CaCl_2,_ 0.1% glycerol, 0.01% LMNG, and 0.001% CHS. The solution was then applied to the protein C antibody affinity resin again to capture uncleaved material. The flowthrough was concentrated and purified by size-exclusion chromatography on a Superdex S200 increase column (Cytiva) equilibrated in 20 mM HEPES pH 7.5, 250 mM NaCl, 0.1% glycerol, 0.01% LMNG, and 0.001% CHS. Peak fractions were pooled, concentrated to 50 mg/ml, aliquoted, flash-frozen in liquid nitrogen, and stored at −80°C until use. Protein purity was assessed by SDS-PAGE.

### Crystallography and data collection

Purified σ_2_ receptor was incorporated into lipidic cubic phase (LCP) by combining the protein with a 10:1 (w:w) mixture of monoolein (Hampton Research) and cholesterol (Sigma Aldrich). Reconstitution was performed at a 1.5:1.0 lipid:protein mass ratio using the coupled syringe method^81^. Each sample was mixed 150 times to ensure formation of a homogeneous phase. Using a Gryphon LCP robot (Art Robbins Instruments), the LCP was dispensed as 30-50 nl drops onto LCP sandwich set (Hampton Research) glass plate and overlaid with 1 μl precipitant solution. ZINCkH000002xsk4-bound crystals grew in 24% PEG 300, 100 mM MES pH 6, 600 mM ammonium phosphate, 100 mM EDTA, and 1 µM ligand. Crystals were harvested with mesh loops (MiTeGen) and stored in liquid nitrogen until data collection.

Diffraction data were collected at Advanced Photon Source beamline 23ID-D using a 10-μm beam. Images were recorded with 0.2° oscillations at a wavelength of 0.47686 Å. The final dataset was generated by merging two datasets collected from two independent crystals.

### Data reduction and refinement

Diffraction images were integrated, scaled, and merged using DIALS^82^. Data collection and processing statistics are reported in **Table 1**. The ZINCkH000002xsk4-bound σ_2_ receptor structure was determined by molecular replacement using PDB ID: 7M95. Matthews probability analysis indicated the presence of one receptor molecule in the asymmetric unit, and a single copy of the search model was placed with Phaser^83^.

The molecular replacement solution was refined through iterative rounds of model adjustment and refinement. Manual model rebuilding was performed in Coot^84^, and refinement was carried out in PHENIX^85^. Final refinement statistics are summarized in **Table 1**.

## Data availability

The compounds docked in this study are freely available from ZINC22 databases, https://cartblanche22.docking.org. Computational predictions and results of top-ranked AF3 and DOCK3 poses will be released upon publication. The coordinates and density map for ZINCkH000002xsk4-bound σ_2_ have been deposited in the PDB with an accession code 36EW.

## Code availability

DOCK3.8 code is freely available on [GitHub](https://github.com/docking-org/dock3). The interaction profiler LUNA is freely available at GitHub. The workflow and scripts for curating and benchmarking AF3 on out-of-sample dataset is freely available at [GitHub](https://github.com/Lyu-Lab/benchmarking-af3).

## Acknowledgements

This work was supported by Rockefeller University start-up funds (J.L.), Irma T. Hirschl/Monique Weill-Caulier Trusts (J.L.), the Stavros Niarchos Foundation (SNF) as part of its grant to the SNF Institute for Global Infectious Disease Research at The Rockefeller University (J.L.), and Yale University start-up funds (A.A.). This material is based upon work supported by the National Science Foundation Graduate Research Fellowship Program under Grant No. DGE-2439608 (A.D.). Any opinions, findings, and conclusions or recommendations expressed in this material are those of the authors and do not necessarily reflect the views of the National Science Foundation. This work used the Delta system at the National Center for Supercomputing Applications through allocation BIO250030 provided to author A.D. from the Advanced Cyberinfrastructure Coordination Ecosystem: Services & Support (ACCESS) program, which is supported by U.S. National Science Foundation grants #2138259, #2138286, #2138307, #2137603, #2138296. This work also used computational resources from the Rockefeller University High Performance Computing Resource Center, RRID: SCR_025889. J.L. is supported by Searle Scholars Program. We thank OpenEye Software for the use of Omega and ROCS. We thank Dr. Anum Glasgow for reading this work.

