## Supplemental materials for "AlphaFold3 for Structure-guided Ligand Discovery"

† These authors contributed equally

* Corresponding authors

### Non-biological ligands

As stated by Krishna et al., many of the ligands found in the PDB are non-biological.[1] In other words, these are ligands that are not relevant in a biological setting, but rather are solvents or other crystal additives used to aid in protein structure determination. We adopted the same list of 3-letter component identifiers as those used by Krishna et al., and also added some of our own ligands after further manual inspection.

Table S1: A list of non-biological ligands used for filtering.

```
'NUC', 'ZN', 'CA', 'MG', 'III', 'MN', 'FE', 'CU', 'SF4', 'FE2',
'CO', 'FES', 'GOL', 'NA', 'CL', 'K', 'CU1', 'GOL', 'XE', 'NO2',
'EDO', 'NI', 'BR', 'CD', 'O', 'CS', 'NO', 'TL', 'HG', 'UNL',
'KR', 'SR', 'RB', 'F', 'AG', 'AR', 'U', 'AU', 'MO', 'SE', 'GD',
'YB', 'VX', 'SM', 'LI', 'RE', 'N', 'W', 'OS', 'HO', 'PI',
'EDO', 'PG4', 'OGA', 'SO4', 'HEZ', 'FEO', 'CL', 'DMS', 'ACT',
'MPD', 'GOL', 'NH2', 'CUA', 'SIW', 'PGW', 'IOD', 'BR', '3NI',
'ZRW', '78M', 'UNX', 'MES', 'CCN', 'PO4', 'PEG'
```

### AF3 Input Examples

Users can run AlphaFold3 with many customizations. We provide three examples of input files we used for structure predictions: with template, without template, and ligand-only.

**With template:**

```
{
    "name": "...",
    "sequences": [
        {
            "protein": {
                "id": "A",
                "sequence": "...",
                "unpairedMsa": "",
                "pairedMsa": "",
                "templates": [
                    {
                        "mmcifPath": "/path/to/template.cif",
                        "queryIndices": [
                            17,
                            18,
                            19,
                            ...
                        ],
                        "templateIndices": [
                            0,
                            1,
                            2,
                            ...
                        ]
                    }
                ]
            }
        },
        {
            "ligand": {
                "id": "B",
                "smiles": "..."
            }
        }
    ],
    "modelSeeds": [4, 9, 8, 31, 20],
    "dialect": "alphafold3",
    "version": 2
}
```

**Without template:**

```
{
  "name": "...",
```

```
  "sequences": [
    {
      "protein": {
        "id": "A",
        "sequence": "..."
      }
    },
    {
      "ligand": {
        "id": "B",
        "smiles": "..."
      }
    }
  ],
  "modelSeeds": [4, 9, 8, 31, 20],
  "dialect": "alphafold3",
  "version": 2
}
```

**Ligand-Only:**

```
{
  "name": "ATP",
  "sequences": [
    {
      "ligand": {
        "id": "B",
        "smiles":
"c1nc(c2c(n1)n(cn2)C3C(C(C(O3)COP(=O)(O)OP(=O)(O)OP(=O)(O)O)O)O)N"
      }
    }
  ],
  "modelSeeds": [4, 9, 8, 31, 20],
  "dialect": "alphafold3",
  "version": 1
}
```

### Compounds from the $\sigma_2$ prospective screen

**Table S2**: All experimentally tested compounds (222) from the AF3 screen (114) and DOCK3 screen (108). AF3 confidence metrics (ipTM and mPAE), DOCK3 score, and fraction [$^3$H]-DTG bound.

| ipTM | mPAE | DOCK3 score (kcal/mol) | Fraction [3H]-DTG bound | AF3/ DOCK3 | Ki (nM) |
|---|---|---|---|---|---|
| 0.92 | 2.21 | -39.87 | 1.18 | AF3 | |
| 0.92 | 1.65 | -26.81 | 1.15 | AF3 | |
| 0.94 | 1.36 | -26.96 | 1.09 | AF3 | |
| 0.92 | 1.76 | -18.27 | 1.08 | AF3 | |
| 0.94 | 1.55 | -39.08 | 1.08 | AF3 | |
| 0.94 | 1.61 | -25.2 | 1.05 | AF3 | |
| 0.93 | 1.93 | -30.73 | 1.05 | AF3 | |
| 0.94 | 1.79 | -48.62 | 1.04 | AF3 | |
| 0.94 | 1.5 | -30.08 | 1.04 | AF3 | |
| 0.93 | 1.78 | -42.85 | 1.03 | AF3 | |
| 0.92 | 2.1 | -29.45 | 1.02 | AF3 | |
| 0.93 | 1.58 | -47.16 | 1.01 | AF3 | |
| 0.92 | 1.76 | 1000 | 1.01 | AF3 | |
| 0.93 | 1.92 | 1000 | 1.01 | AF3 | |
| 0.93 | 1.63 | -44.57 | 0.99 | AF3 | |
| 0.93 | 1.93 | -41.44 | 0.99 | AF3 | |
| 0.93 | 1.84 | -35.09 | 0.99 | AF3 | |
| 0.92 | 1.64 | -22.93 | 0.97 | AF3 | |
| 0.92 | 2 | -39.9 | 0.97 | AF3 | |
| 0.92 | 2.18 | -47.44 | 0.96 | AF3 | |

| | | | | |
|---|---|---|---|---|
| 0.92 | 1.88 | 1000 | 0.96 | AF3 |
| 0.93 | 1.58 | -43.14 | 0.95 | AF3 |
| 0.93 | 1.56 | -36.02 | 0.95 | AF3 |
| 0.93 | 1.65 | -37.03 | 0.95 | AF3 |
| 0.93 | 1.67 | -40.78 | 0.95 | AF3 |
| 0.92 | 1.99 | -38.09 | 0.94 | AF3 |
| 0.92 | 1.99 | 1000 | 0.94 | AF3 |
| 0.93 | 1.76 | -41.75 | 0.94 | AF3 |
| 0.94 | 1.55 | -17.67 | 0.94 | AF3 |
| 0.92 | 1.81 | -39.51 | 0.93 | AF3 |
| 0.93 | 1.72 | -46.14 | 0.93 | AF3 |
| 0.93 | 2.05 | -38.63 | 0.93 | AF3 |
| 0.93 | 1.63 | -27.25 | 0.93 | AF3 |
| 0.93 | 1.51 | -40.64 | 0.93 | AF3 |
| 0.92 | 1.81 | 1000 | 0.93 | AF3 |
| 0.93 | 1.49 | -28.85 | 0.92 | AF3 |
| 0.93 | 1.66 | 12.06 | 0.92 | AF3 |
| 0.93 | 1.74 | -36.52 | 0.92 | AF3 |
| 0.94 | 1.81 | -52.85 | 0.91 | AF3 |
| 0.94 | 1.41 | -35.1 | 0.91 | AF3 |
| 0.92 | 2.09 | -37.9 | 0.91 | AF3 |
| 0.94 | 1.78 | -50.46 | 0.91 | AF3 |
| 0.93 | 2.11 | -21.45 | 0.91 | AF3 |
| 0.92 | 1.66 | 1000 | 0.91 | AF3 |
| 0.93 | 2.03 | -44.06 | 0.91 | AF3 |
| 0.93 | 1.31 | 1000 | 0.91 | AF3 |
| 0.92 | 1.74 | -43.33 | 0.91 | AF3 |
| 0.93 | 1.65 | -39.25 | 0.90 | AF3 |

| | | | | |
|---|---|---|---|---|
| 0.94 | 1.41 | 1000 | 0.90 | AF3 |
| 0.93 | 1.76 | -38.54 | 0.90 | AF3 |
| 0.93 | 1.91 | -43.61 | 0.90 | AF3 |
| 0.94 | 1.38 | 1000 | 0.90 | AF3 |
| 0.92 | 1.62 | 1000 | 0.90 | AF3 |
| 0.93 | 1.71 | -37.03 | 0.90 | AF3 |
| 0.93 | 1.99 | -45.26 | 0.90 | AF3 |
| 0.92 | 1.96 | -42.31 | 0.89 | AF3 |
| 0.93 | 1.7 | -42.37 | 0.89 | AF3 |
| 0.92 | 1.99 | -39.18 | 0.88 | AF3 |
| 0.94 | 1.34 | -41.13 | 0.88 | AF3 |
| 0.92 | 1.92 | 1000 | 0.88 | AF3 |
| 0.92 | 1.6 | -30.5 | 0.88 | AF3 |
| 0.93 | 1.5 | -41.16 | 0.87 | AF3 |
| 0.92 | 1.98 | -44.53 | 0.87 | AF3 |
| 0.93 | 1.96 | -43.61 | 0.86 | AF3 |
| 0.93 | 1.59 | -28.47 | 0.86 | AF3 |
| 0.93 | 2.01 | -45.95 | 0.85 | AF3 |
| 0.93 | 1.78 | -36.97 | 0.85 | AF3 |
| 0.93 | 1.85 | -38.46 | 0.85 | AF3 |
| 0.93 | 1.31 | -39.64 | 0.85 | AF3 |
| 0.93 | 1.68 | -33.56 | 0.84 | AF3 |
| 0.93 | 1.78 | -39.63 | 0.84 | AF3 |
| 0.93 | 2.14 | -33.2 | 0.83 | AF3 |
| 0.94 | 1.7 | -43.55 | 0.83 | AF3 |
| 0.94 | 1.42 | -45.96 | 0.82 | AF3 |
| 0.93 | 1.7 | -40.91 | 0.82 | AF3 |
| 0.94 | 1.22 | -46.62 | 0.80 | AF3 |

| | | | | |
|---|---|---|---|---|
| 0.93 | 1.81 | -46.26 | 0.79 | AF3 |
| 0.92 | 1.57 | -32.17 | 0.78 | AF3 |
| 0.93 | 1.5 | -35.57 | 0.77 | AF3 |
| 0.92 | 2 | -36.85 | 0.77 | AF3 |
| 0.93 | 1.65 | -36.29 | 0.76 | AF3 |
| 0.93 | 1.33 | -30.34 | 0.75 | AF3 |
| 0.95 | 1.16 | -50.3 | 0.75 | AF3 |
| 0.93 | 1.57 | -37.05 | 0.73 | AF3 |
| 0.94 | 1.64 | -30.8 | 0.73 | AF3 |
| 0.93 | 1.88 | -38.57 | 0.72 | AF3 |
| 0.94 | 1.5 | -49.96 | 0.72 | AF3 |
| 0.93 | 1.93 | -46.49 | 0.71 | AF3 |
| 0.92 | 2.15 | -37.14 | 0.71 | AF3 |
| 0.95 | 1.04 | -41.79 | 0.70 | AF3 |
| 0.92 | 1.73 | -39.35 | 0.70 | AF3 |
| 0.92 | 1.92 | -33.62 | 0.66 | AF3 |
| 0.93 | 1.25 | -32.97 | 0.64 | AF3 |
| 0.93 | 1.22 | -42.96 | 0.63 | AF3 |
| 0.93 | 1.44 | -49.52 | 0.62 | AF3 |
| 0.92 | 1.92 | -53.61 | 0.62 | AF3 |
| 0.93 | 1.68 | -44.49 | 0.61 | AF3 |
| 0.93 | 1.56 | 1000 | 0.60 | AF3 |
| 0.92 | 1.99 | 1000 | 0.57 | AF3 |
| 0.94 | 1.64 | 1000 | 0.49 | AF3 |
| 0.94 | 1.4 | -45.46 | 0.39 | AF3 |
| 0.93 | 1.65 | -47.19 | 0.37 | AF3 |
| 0.95 | 1.44 | -41.89 | 0.34 | AF3 |
| 0.94 | 1.43 | -44.4 | 0.33 | AF3 |
| 0.94 | 1.41 | 1000 | 0.28 | AF3 |

| | | | | | |
|---|---|---|---|---|---|
| 0.92 | 2.13 | -36.36 | 0.25 | AF3 | |
| 0.94 | 1.35 | -39.36 | 0.18 | AF3 | |
| 0.93 | 1.74 | -48.88 | 0.18 | AF3 | |
| 0.94 | 1.55 | 1000 | 0.13 | AF3 | |
| 0.92 | 2.33 | -38.01 | 0.11 | AF3 | 131 |
| 0.94 | 1.68 | -40.48 | 0.08 | AF3 | 107 |
| 0.93 | 1.7 | -47.3 | 0.07 | AF3 | 13 |
| 0.93 | 1.77 | -40.06 | 0.05 | AF3 | 38 |
| 0.93 | 1.95 | -46.9 | 0.02 | AF3 | 42 |
| 0.86 | 2.66 | -52.26 | 1.20 | DOCK3 | |
| 0.8 | 4.3 | -54.69 | 1.12 | DOCK3 | |
| 0.83 | 4.22 | -57.58 | 1.11 | DOCK3 | |
| 0.91 | 2.38 | -53.56 | 1.11 | DOCK3 | |
| 0.82 | 3.51 | -56.4 | 1.06 | DOCK3 | |
| 0.92 | 1.88 | -53.84 | 1.05 | DOCK3 | |
| 0.8 | 4.11 | -57.14 | 1.02 | DOCK3 | |
| 0.89 | 2.49 | -57.37 | 1.01 | DOCK3 | |
| 0.83 | 3.63 | -54.45 | 1.01 | DOCK3 | |
| 0.8 | 3.84 | -58.5 | 1.00 | DOCK3 | |
| 0.9 | 2.55 | -57.01 | 1.00 | DOCK3 | |
| 0.82 | 3.74 | -53.51 | 0.98 | DOCK3 | |
| 0.87 | 3.52 | -55 | 0.98 | DOCK3 | |
| 0.83 | 3.73 | -54.53 | 0.96 | DOCK3 | |
| 0.89 | 2.69 | -52.73 | 0.96 | DOCK3 | |
| 0.92 | 1.88 | -52.64 | 0.96 | DOCK3 | |
| 0.77 | 4.49 | -53.14 | 0.96 | DOCK3 | |
| 0.77 | 4.4 | -56.75 | 0.96 | DOCK3 | |
| 0.93 | 1.44 | -53.21 | 0.96 | DOCK3 | |

| | | | | |
|---|---|---|---|---|
| 0.81 | 3.61 | -56.82 | 0.94 | DOCK3 |
| 0.86 | 2.94 | -56.57 | 0.93 | DOCK3 |
| 0.82 | 3.95 | -53.94 | 0.93 | DOCK3 |
| 0.78 | 4.43 | -54.73 | 0.93 | DOCK3 |
| 0.83 | 3.71 | -52.96 | 0.92 | DOCK3 |
| 0.87 | 2.98 | -56.48 | 0.91 | DOCK3 |
| 0.8 | 3.69 | -53.23 | 0.91 | DOCK3 |
| 0.81 | 4.19 | -53.86 | 0.91 | DOCK3 |
| 0.81 | 4.08 | -53.66 | 0.90 | DOCK3 |
| 0.82 | 3.62 | -55.27 | 0.90 | DOCK3 |
| 0.81 | 3.5 | -55.07 | 0.89 | DOCK3 |
| 0.8 | 3.43 | -53.42 | 0.89 | DOCK3 |
| 0.9 | 2.19 | -58.74 | 0.89 | DOCK3 |
| 0.85 | 3.32 | -58.77 | 0.89 | DOCK3 |
| 0.8 | 3.46 | -53.36 | 0.88 | DOCK3 |
| 0.81 | 3.92 | -52.73 | 0.88 | DOCK3 |
| 0.88 | 2.57 | -53.71 | 0.86 | DOCK3 |
| 0.76 | 4.42 | -56.85 | 0.85 | DOCK3 |
| 0.89 | 2.64 | -52.29 | 0.84 | DOCK3 |
| 0.89 | 2.45 | -55.07 | 0.83 | DOCK3 |
| 0.85 | 3.31 | -53.85 | 0.83 | DOCK3 |
| 0.88 | 2.81 | -54.47 | 0.83 | DOCK3 |
| 0.86 | 3.3 | -54.12 | 0.83 | DOCK3 |
| 0.87 | 2.7 | -52.85 | 0.83 | DOCK3 |
| 0.8 | 4.18 | -54.32 | 0.83 | DOCK3 |
| 0.87 | 2.87 | -58.74 | 0.82 | DOCK3 |
| 0.78 | 4.57 | -53.43 | 0.82 | DOCK3 |
| 0.81 | 4.32 | -55.22 | 0.81 | DOCK3 |

| | | | | |
|---|---|---|---|---|
| 0.85 | 2.9 | -58.27 | 0.81 | DOCK3 |
| 0.85 | 2.87 | -53.15 | 0.81 | DOCK3 |
| 0.9 | 2.66 | -53.23 | 0.81 | DOCK3 |
| 0.83 | 3.89 | -54.34 | 0.78 | DOCK3 |
| 0.87 | 2.68 | -52.74 | 0.78 | DOCK3 |
| 0.85 | 3.43 | -54.73 | 0.78 | DOCK3 |
| 0.84 | 3.38 | -62.55 | 0.77 | DOCK3 |
| 0.75 | 4.39 | -53.5 | 0.77 | DOCK3 |
| 0.8 | 4.35 | -52.87 | 0.76 | DOCK3 |
| 0.8 | 3.91 | -54.51 | 0.76 | DOCK3 |
| 0.85 | 3.15 | -54.2 | 0.75 | DOCK3 |
| 0.81 | 3.91 | -52.81 | 0.75 | DOCK3 |
| 0.88 | 2.99 | -52.63 | 0.73 | DOCK3 |
| 0.84 | 3.89 | -54.38 | 0.72 | DOCK3 |
| 0.79 | 3.83 | -54.01 | 0.71 | DOCK3 |
| 0.78 | 4.31 | -52.95 | 0.71 | DOCK3 |
| 0.8 | 4.16 | -54.3 | 0.70 | DOCK3 |
| 0.79 | 4.09 | -54.28 | 0.69 | DOCK3 |
| 0.77 | 4.21 | -52.74 | 0.68 | DOCK3 |
| 0.86 | 3.57 | -52.28 | 0.64 | DOCK3 |
| 0.87 | 2.87 | -57.76 | 0.64 | DOCK3 |
| 0.81 | 3.9 | -55.9 | 0.63 | DOCK3 |
| 0.86 | 3.48 | -56.7 | 0.62 | DOCK3 |
| 0.87 | 3.44 | -54.62 | 0.60 | DOCK3 |
| 0.91 | 2.19 | -56.56 | 0.60 | DOCK3 |
| 0.81 | 4.23 | -60.54 | 0.59 | DOCK3 |
| 0.79 | 4.18 | -52.91 | 0.58 | DOCK3 |
| 0.81 | 3.98 | -53.44 | 0.56 | DOCK3 |
| 0.82 | 3.77 | -52.79 | 0.56 | DOCK3 |

| | | | | | |
|---|---|---|---|---|---|
| 0.76 | 4.84 | -55.41 | 0.55 | DOCK3 | |
| 0.84 | 3.42 | -54.65 | 0.54 | DOCK3 | |
| 0.85 | 3.16 | -52.97 | 0.54 | DOCK3 | |
| 0.86 | 2.91 | -53.04 | 0.50 | DOCK3 | |
| 0.83 | 3.91 | -55.88 | 0.48 | DOCK3 | |
| 0.83 | 4 | -60.36 | 0.43 | DOCK3 | |
| 0.9 | 2.36 | -52.93 | 0.41 | DOCK3 | |
| 0.81 | 4.42 | -53.99 | 0.40 | DOCK3 | |
| 0.77 | 4.79 | -57.25 | 0.39 | DOCK3 | |
| 0.89 | 2.83 | -54.31 | 0.37 | DOCK3 | |
| 0.83 | 3.83 | -53.31 | 0.37 | DOCK3 | |
| 0.76 | 4.87 | -52.96 | 0.34 | DOCK3 | |
| 0.91 | 2.26 | -60.4 | 0.31 | DOCK3 | |
| 0.79 | 4.54 | -55.85 | 0.28 | DOCK3 | |
| 0.82 | 3.85 | -55.06 | 0.25 | DOCK3 | |
| 0.81 | 3.46 | -60.3 | 0.24 | DOCK3 | |
| 0.85 | 3.65 | -57.67 | 0.23 | DOCK3 | |
| 0.87 | 3.07 | -53.54 | 0.15 | DOCK3 | |
| 0.9 | 2.36 | -54.55 | 0.15 | DOCK3 | |
| 0.77 | 4.49 | -55.19 | 0.14 | DOCK3 | |
| 0.8 | 3.64 | -54.57 | 0.12 | DOCK3 | |
| 0.84 | 3.87 | -56.95 | 0.11 | DOCK3 | |
| 0.84 | 3.73 | -53.99 | 0.11 | DOCK3 | |
| 0.74 | 5.47 | -52.72 | 0.08 | DOCK3 | |
| 0.8 | 3.93 | -52.63 | 0.07 | DOCK3 | |
| 0.85 | 3.54 | -53.97 | 0.06 | DOCK3 | 199 |
| 0.78 | 4.04 | -54.14 | 0.05 | DOCK3 | 113 |

| | | | | | |
|---|---|---|---|---|---|
| 0.88 | 3.14 | -53.37 | 0.04 | DOCK3 | 39 |
| 0.77 | 4.8 | -54.73 | 0.02 | DOCK3 | 43 |
| 0.86 | 2.97 | -53.37 | 0.00 | DOCK3 | 66 |
| 0.85 | 3.19 | -54.31 | 0.88 | DOCK3 | |
| 0.86 | 3.22 | -52.2 | 0.97 | DOCK3 | |
